# Hypomodified tRNA in evolutionarily distant yeasts can trigger rapid tRNA decay to activate the general amino acid control response, but with different consequences

**DOI:** 10.1101/2020.05.29.123083

**Authors:** Thareendra De Zoysa, Eric M. Phizicky

## Abstract

All tRNAs are extensively modified, and modification deficiency often results in growth defects in the budding yeast *Saccharomyces cerevisiae* and neurological or other disorders in humans. In *S. cerevisiae*, lack of any of several tRNA body modifications results in rapid tRNA decay (RTD) of certain mature tRNAs by the 5’-3’ exonucleases Rat1 and Xrn1. As tRNA quality control decay mechanisms are not extensively studied in other eukaryotes, we studied *trm8Δ* mutants in the evolutionarily distant fission yeast *Schizosaccharomyces pombe*, which lack 7-methylguanosine at G_46_ of tRNAs. We report here that *S. pombe trm8Δ* mutants are temperature sensitive primarily due to decay of tRNA^Tyr(GUA)^ and that spontaneous mutations in the *RAT1* ortholog *dhp1^+^* restored temperature resistance and prevented tRNA decay, demonstrating conservation of the RTD pathway. We also report for the first time evidence linking the RTD and the general amino acid control (GAAC) pathways, which we show in both *S. pombe* and *S. cerevisiae*. In *S. pombe trm8Δ* mutants, spontaneous GAAC mutations restored temperature resistance and tRNA levels, and the temperature sensitivity of *trm8Δ* mutants was precisely linked to GAAC activation due to tRNA^Tyr(GUA)^ decay. Similarly, in the well-studied *S. cerevisiae trm8Δ trm4Δ* RTD mutant, temperature sensitivity was closely linked to GAAC activation due to tRNA^Val(AAC)^ decay; however, in *S. cerevisiae*, GAAC mutations increased tRNA decay and enhanced temperature sensitivity. Thus, these results demonstrate a conserved GAAC activation coincident with RTD in *S. pombe* and *S. cerevisiae*, but an opposite impact of the GAAC response in the two organisms. We speculate that the RTD pathway and its regulation of the GAAC pathway is widely conserved in eukaryotes, extending to other mutants affecting tRNA body modifications.

**Author Summary:** tRNA modifications are highly conserved and their lack frequently results in growth defects in the yeast *Saccharomyces cerevisiae* and neuorological disorders in humans. *S. cerevsiaie* has two tRNA quality control decay pathways that sense tRNAs lacking modifications in the main tRNA body. One of these, the rapid tRNA decay (RTD) pathway, targets mature tRNAs for 5’-3’ exonucleolytic decay by Rat1 and Xrn1. It is unknown if RTD is conserved in eukaryotes, and if it might explain phenotypes associated with body modification defects. Here we focus on *trm8Δ* mutants, lacking m^7^G_46_, in the evolutionarily distant yeast *Schizosaccharomyces pombe*. Loss of m^7^G causes temperature sensitivity and RTD in *S. cerevisiae*, microcephalic primordial dwarfism in humans, and defective stem cell renewal in mice. We show that *S. pombe trm8Δ* mutants are temperature sensitive due to tY(GUA) decay by Rat1, implying conservation of RTD among divergent eukaryotes. We also show that the onset of RTD triggers activation of the general amino acid control (GAAC) pathway in both *S. pombe* and *S. cerevisiae*, resulting in exacerbated decay in *S. pombe* and reduced decay in *S. cerevisiae*. We speculate that RTD and its regulation of the GAAC pathway will be widely conserved in eukaryotes including humans.

## Introduction

tRNAs are subject to extensive post-transcriptional modifications that often profoundly affect tRNA function, as lack of modifications often leads to growth defects in the budding yeast *Saccharomyces cerevisiae* and to neurological or mitochondrial disorders in humans (1–5). Many tRNA modifications in the anticodon loop are important for decoding fidelity, reading frame maintenance, and sometimes charging efficiency (6–15). By contrast, modifications in the tRNA body, the region outside the anticodon loop, are often important for folding and stability (16–18), resulting in substantial growth defects. In *S. cerevisiae*, deletion of *TRM6* or *TRM61* is lethal, associated with lack of 1-methyladenosine at A_58_ (m^1^A_58_) (19), whereas deletion of *TAN1*, *TRM1*, or *TRM8* (or *TRM82*) results in temperature sensitivity associated with lack of 4-acetylcytidine at C_12_ (ac^4^C_12_), N_2_,N_2_- dimethylguanosine at G_26_ (m^2,2^G_26_), or 7-methylguanosine at G_46_ (m^7^G_46_) respectively (20–22). Similarly, human neurological disorders are linked to mutations in *TRMT10A*, associated with reduced 1-methylguanosine at G_9_ (m^1^G_9_) (23, 24), *TRMT1* (m^2,2^G_26_) (25–28), *WDR4* (m^7^G_46_) (29–31) and *NSUN2*, associated with reduced 5-methylcytidine (m^5^C) at C_48-50_, as well as at C_34_ and C_40_ (32–34).

In *S. cerevisiae*, lack of any of several tRNA body modifications leads to decay of a subset of the corresponding hypomodified tRNAs, mediated by either of two tRNA quality control pathways, each acting on different hypomodified tRNAs and at different stages of tRNA biogenesis. First, the nuclear surveillance pathway targets pre-tRNA_i_^Met^ lacking m^1^A, acting through the the nuclear exosome and the TRAMP complex to degrade the pre-tRNA from the 3’ end (17, 35–37). The nuclear surveillance pathway also targets a large portion of wild type (WT) pre-tRNAs shortly after transcription, ascribed to errors in folding of the nascent tRNA or to mutations arising during transcription (38). Second, the rapid tRNA decay (RTD) pathway targets certain mature tRNAs lacking m^7^G_46_, m^2,2^G_26_, or ac^4^C_12_, using the 5’-3’ exonucleases Rat1 in the nucleus and Xrn1 in the cytoplasm (18, 21, 22, 39, 40). RTD is inhibited by a *met22Δ* mutation (22, 39, 41, 42) due to accumulation of the Met22 substrate adenosine 3’, 5’ bisphosphate (pAp) (43, 44), which binds the active site of Xrn1 and presumably Rat1 (45). The RTD pathway also targets fully modified tRNAs with destabilizing mutations in the stems, particularly the acceptor and T-stem, which expose the 5’ end (40–42). The hypomodified tRNAs targeted by the RTD pathway also expose the 5’ end, ascribed to destabilization of the tertiary fold (40).

There is limited evidence documenting tRNA quality control decay pathways that act on hypomodified tRNAs in other eukaryotes. A mouse embryonic stem cell line with a knockout of *METTL1* (ortholog of *S. cerevisiae TRM8*) had undetectable m^7^G in its tRNA substrates and reduced levels of several METTL1 substrate tRNAs (46). Similarly, knockdown of *METTL1* and *NSUN2* (homolog of *S. cerevisiae TRM4*) in HeLa cells led to reduced levels of tRNA^Val(AAC)^ (tV(AAC)) at 43°C in the presence of 5-fluorouracil (5-FU) (47), a known inhibitor of pseudouridine synthases and 5-methyluridine methyltransferase (48–50). However, in both of these cases, the underlying mechanism is not known. It was also shown that WT mature tRNAi^Met^ was subject to decay by Xrn1 and Rat1 after 43°C heat shock in HeLa cells, although there was no change in the modification pattern *in vivo* or in physical stability of the tRNA *in vitro* caused by this temperature shift (51).

The goal of the work described here is to determine if and to what extent tRNA quality control decay pathways are linked to hypomodified tRNAs in eukaryotes other than *S. cerevisiae*. To address this issue, we have studied the biology of the tRNA m^7^G_46_ methyltransferase Trm8 in the fission yeast *Schizosaccharomyces pombe,* which diverged from *S. cerevisiae ∼* 600 million years ago (52).

We chose to study *S.pombe* Trm8 because *S. cerevisiae trm8Δ* mutants were known to trigger decay by the RTD pathway. *S. cerevisiae* Trm8 forms a complex with Trm82 that is required for formation of m^7^G_46_ in eukaryotic tRNAs (53, 54). *S. cerevisiae trm8Δ* and *trm82Δ* mutants are each temperature sensitive (20), and *trm8Δ* or *trm82Δ* mutants also lacking any of several other body modifications had enhanced temperature sensitivity (18). Moreover, the temperature sensitivity of *trm8Δ* mutants was suppressed by a *met22Δ* mutation and was associated with decay of tV(AAC) (22), and the temperature sensitivity of *trm8Δ trm4Δ* mutants (lacking both m^7^G and m^5^C) was shown explicitly to be due to RTD of tV(AAC) (18, 39). Furthermore, Trm8 biology in mammalian cells has additional dimensions of complexity. The human *TRM82* ortholog *WDR4* was associated with reduced tRNA m^7^G modification and a distinct form of microcephalic primordial dwarfism (29); *METTL1* or *WDR4* knock out mouse embryonic stem cells showed defects in self renewal and differentiation (46); and METTL1 was also responsible for m^7^G modification of mammalian miRNAs and mRNAs (55, 56). This evidence emphasizes that Trm8/Trm82 (METTL1/WDR4) and/or its m^7^G modification product is important in *S. cerevisiae* and mammals, although the reasons are not yet known beyond *S. cerevisiae*.

We find here that *S. pombe trm8Δ* mutants have a temperature sensitive growth defect due primarily to decay of tY(GUA) and to some extent tP(AGG) by the Rat1 ortholog Dhp1, demonstrating that a major component of the RTD pathway is conserved between *S. pombe* and *S. cerevisiae.* We also find an unexpected connection between the RTD pathway and the general amino acid control (GAAC) pathway in both *S. pombe* and *S. cerevisiae*. In both *S. pombe trm8Δ* mutants and *S. cerevisiae trm8Δ trm4Δ* mutants, the temperature sensitivity coincides with the onset of tRNA decay, which in turn triggers the GAAC activation, presumably due to the increased stress by the tRNA decay. However, in *Sp trm8Δ* mutants, GAAC activation is deleterious to growth, as mutations in the GAAC pathway restore growth and tRNA levels, whereas in *S. cerevisiae trm8Δ trm4Δ* mutants, GAAC pathway activation is beneficial, as GAAC mutations exacerbate the growth defect and accelerate tRNA decay. Thus, our results demonstrate a conserved GAAC response associated with tRNA decay but opposite effects on cell physiology in the two organisms. These findings suggest the widespread conservation of the RTD pathway in eukaryotes, and the widespread conservation of the GAAC pathway in monitoring overall tRNA quality.

## Results

### The *S. pombe trm8Δ* mutants lack m^7^G in tRNAs and are temperature sensitive

As Trm8 is the catalytic subunit of the Trm8-Trm82 complex (20), we anticipated that tRNAs from *S. pombe trm8Δ* mutants would lack m^7^G. We purified tY(GUA) and tF(GAA), which had each been previously shown to have m^7^G_46_ (57, 58), and then analyzed their nucleosides by HPLC analysis. Purified tY(GUA) from *S. pombe trm8Δ* mutants had no detectable m^7^G levels (less than 0.03 moles/mole), compared to near stoichiometric levels in tY(GUA) from WT cells (0.93 +/- 0.22 moles/mole), whereas levels of each of three other analyzed modifications (pseudouridine (Ψ), m^5^C, and m^1^A) were very similar in *trm8Δ* and WT cells (Fig. 1A). Similarly, purified tF(GAA) from *trm8Δ* mutants had no detectable levels of m^7^G compared to near stoichiometric levels in WT cells, but otherwise WT levels of Ψ, 2’-O-methylcytidine (Cm) and m^2,2^G (Fig. 1A). These results suggest strongly that *S. pombe trm8^+^* is the methyltransferase responsible for m^7^G formation in cytoplasmic tRNAs.

**Fig. 1.**
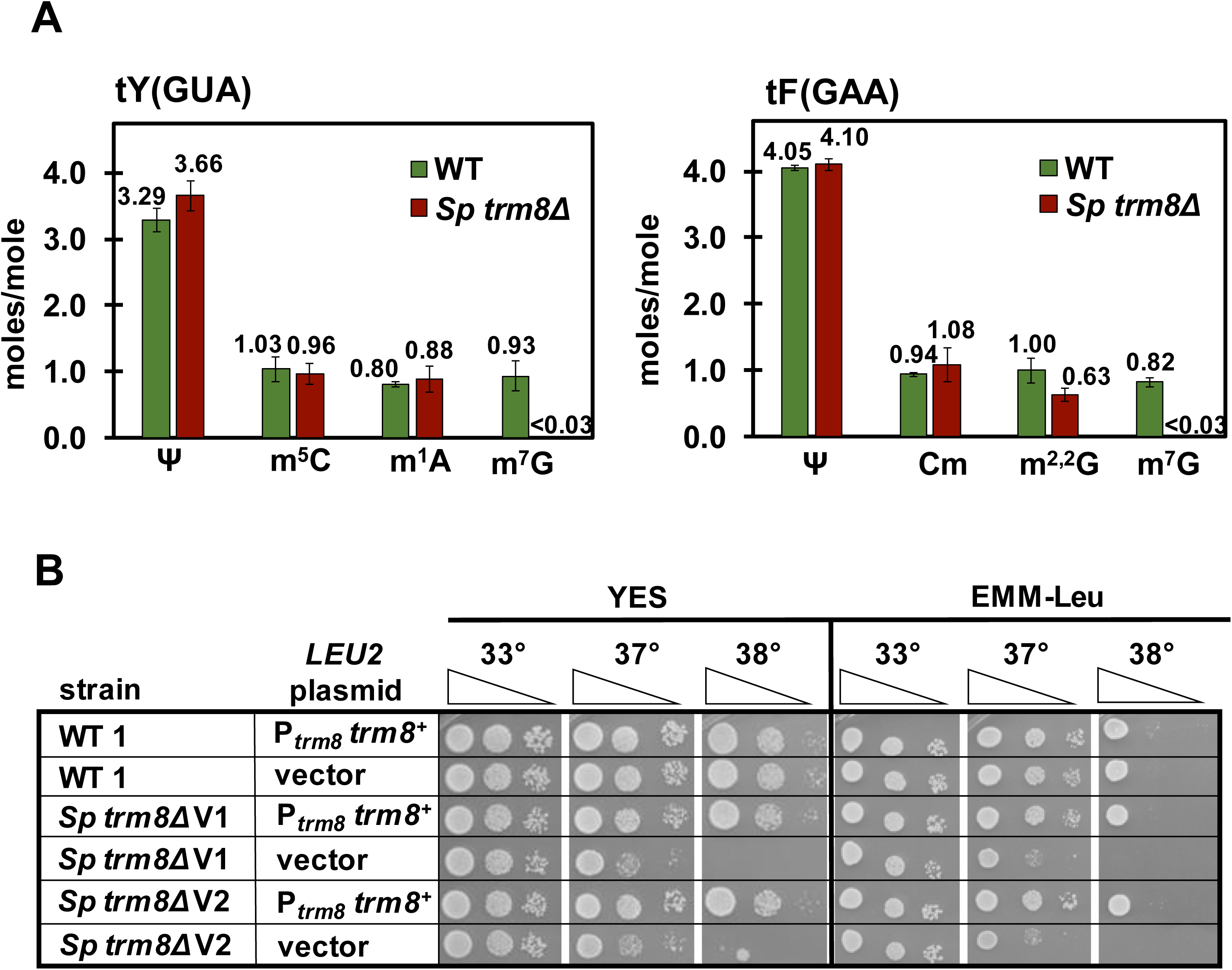
*S. pombe trm8Δ* mutants lacked m^7^G and were temperature sensitive. *(A) trm8Δ* mutants had no detectable m^7^G in their tY(GUA) and tF(GAA). *S. pombe trm8Δ* mutants and WT cells were grown in biological triplicate in YES media at 30°C and tRNAs were purified, digested to nucleosides, and analyzed for modifications by HPLC as described in Materials and Methods. The bar chart depicts average moles/mol values of nucleosides with associated standard deviation; WT, green; *S. pombe* (*Sp*) *trm8Δ,* brown. *(B) trm8Δ* mutants were temperature sensitive due to lack of *trm8^+^.* Strains with plasmids as indicated were grown overnight in EMM-Leu media at 30°C, diluted to OD_600_ ∼ 0.5, serially diluted 10-fold in EMM-Leu, and 2 μL was spotted onto plates containing EMM-Leu or YES media and incubated at 33°C, 37°C, and 38°C. The two independent *trm8Δ* mutants were labeled as *Sp trm8Δ* V1 and V2.

To understand the biology of *S. pombe trm8Δ* mutants, we examined the growth phenotypes of two genetically independent *trm8Δ* mutants. *trm8Δ* mutants were temperature sensitive starting at 37°C on rich (YES) and minimal (EMM) media, and expression of P*_trm8_ trm8^+^* on a plasmid restored WT growth in both media (Fig. 1B). Thus, the temperature sensitivity of *trm8Δ* mutants was due to lack of *trm8^+^*.

### *S. pombe trm8Δ* mutants have reduced levels of tP(AGG) and tY(GUA) at high temperatures

To determine if the temperature sensitivity of *S. pombe trm8Δ* mutants was associated with tRNA decay, we analyzed tRNA levels of *trm8Δ* mutants after an 8 hour temperature shift in YES media from 30°C to 36.5°C, 37.5°C, and 38.5°C, which progressively inhibited growth (Fig. S1). We measured tRNA levels of all 21 tRNAs in the Genomic tRNA Database (59) that had a 5-nt variable loop with a central guanosine residue (Table S1), which is the signature for m^7^G modification (60). We quantified levels of each tRNA at each temperature relative to the levels of that tRNA in WT cells at 30°C, after normalization of each to the levels of the non-Trm8 substrate tG(GCC) at the corresponding temperature. We used tG(GCC) as the standard because the usual standards, 5S and 5.8S RNA, each had temperature-dependent reduction in their levels in *trm8Δ* mutants (Fig S2), as determined relative to input RNA levels. Note that with tG(GCC) as the standard, the levels of another non-Trm8 substrate, tL(UAA), were also unaffected.

Northern analysis showed that *S. pombe trm8Δ* mutants had significantly reduced levels of two of the 21 potential Trm8 substrate tRNAs as the temperature was increased. The levels of tP(AGG) were substantially reduced in *trm8Δ* mutants, from 70% of the levels in WT cells at 30°C, to 50%, 31%, and 18% after temperature shift to 36.5°C, 37.5°C, and 38.5°C respectively, whereas levels of tP(AGG) in WT cells remained constant as temperature increased (Fig. 2A, 2B). As expected, tP(AGG) is indeed a substrate of Trm8, since purified tP(AGG) from *trm8Δ* mutants had undetectable levels of m^7^G, but WT levels of each of three other modifications (Fig. S3). The levels of tY(GUA) were also reduced in *trm8Δ* mutants as temperature increased, albeit to a lesser extent than tP(AGG) levels. Levels of tY(GUA) in *trm8Δ* mutants were about the same as those in WT cells at 30°C (119%), remained essentially unchanged at 36.5°C and 37.5°C (124%, and 98%), but were reduced to 67% at 38.5°C. Levels of tY(GUA) in WT cells were relatively constant at all temperatures. In contrast, none of the 19 other predicted Trm8 substrate tRNAs showed a temperature-dependent reduction in levels (Fig. 2A, 2B, S4, S5). Levels of 15 tRNAs were approximately constant in *trm8Δ* mutants as temperature increased, although the initial levels varied somewhat, and levels of the other four tRNAs (tR(CCU), tM_e_(CAU), tV(CAC), and tK(UUU)) were modestly increased at 38.5°C. Thus, if the temperature sensitivity of *trm8Δ* mutants was due to loss of tRNAs, the likely candidates were tP(AGG) and tY(GUA).

**Fig. 2.**
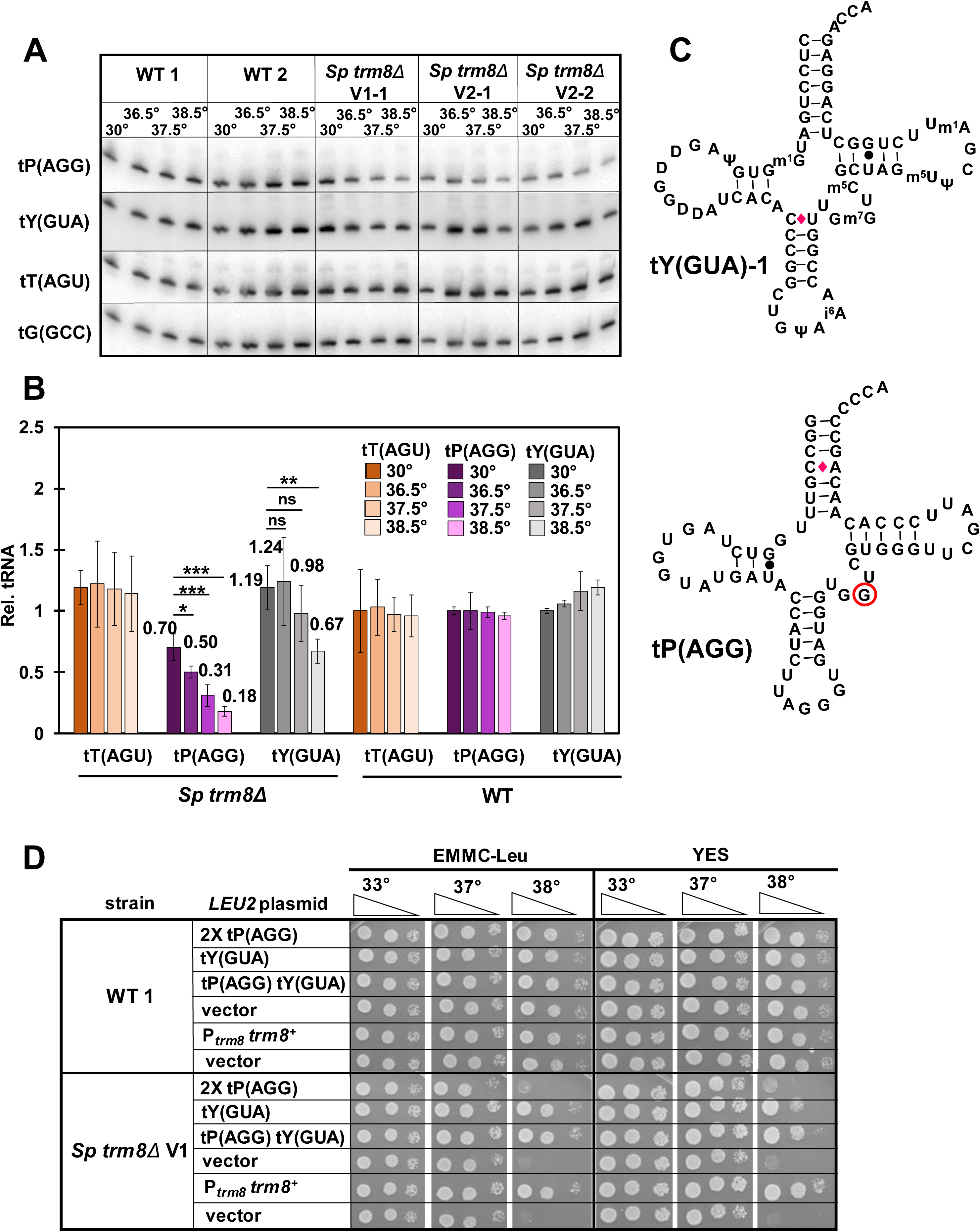
*S. pombe trm8Δ* mutants had reduced levels of tP(AGG) and tY(GUA) at elevated temperatures. *(A)* Northern analysis of Trm8 substrates tP(AGG), tY(GUA), and tT(AGU) in *trm8Δ* and WT cells after shift from 30°C to 36.5°C, 37.5°C, and 38.5°C. Strains were grown in YES media at 30°C, shifted to the indicated temperatures for 8 hours as described in Materials and Methods, and RNA was isolated and analyzed by northern blotting. n=2 for WT cells, n=3 for *S. pombe trm8Δ* mutants. *(B)* Quantification of tP(AGG), tY(GUA), and tT(AGU) levels in WT and *trm8Δ* mutants at different temperatures. The bar chart depicts relative levels of tRNA species at each temperature, relative to their levels in the WT strain at 30°C (each value itself first normalized to levels of the control non-Trm8 substrate tG(GCC)). For each tRNA, lighter shades indicate progressively higher temperatures (30°C, 36.5°C, 37.5°C to 38.5°C) for tT(AGU), brown; tP(AGG), purple; tY(GUA), gray. Standard deviations for each tRNA measurement are indicated. The statistical significance of tRNA levels was evaluated using a two-tailed Student’s t-test assuming equal variance. ns, not significant (p > 0.05); *, p < 0.05; **, p < 0.01; ***, p < 0.001. *(C)* Schematic of the secondary structure of tY(GUA)-1 and tP(AGG). Modifications of tY(GUA) are as annotated. WC base pairs, black lines; GU base pairs, black dots; mismatch C-A or C-U base pairs, red diamonds; presumed m^7^G_46_, red circle. *(D)* Overproduction of tY(GUA), but not tP(AGG), suppressed the temperature sensitive growth defect of *trm8Δ* mutants. Strains with plasmids as indicated were grown overnight in EMMC-Leu media at 30°C and analyzed for growth as in Fig. 1B on the indicated plates.

### The growth defect of *S. pombe trm8Δ* mutants is primarily due to loss of tY(GUA)

To evaluate the cause of the temperature sensitivity of *S. pombe trm8Δ* mutants, we analyzed growth after overexpression of tP(AGG) and/or tY(GUA) on plasmids (Fig. 2C). Surprisingly, on both YES media and EMM complete (EMMC) media, *trm8Δ* mutants expressing tY(GUA) grew almost as well as the *trm8Δ* [P*_trm8_ trm8^+^*] strain or the WT strain at elevated temperatures, whereas *trm8Δ* mutants expressing two tP(AGG) genes on a plasmid had little effect on the temperature sensitivity (Fig. 2D). As expected, northern analysis showed that *trm8Δ* [*leu2^+^* tY(GUA)] strains had substantially more tY(GUA) than the *trm8Δ* [*leu2^+^*] vector control strain at 30°C and 38.5°C (3.2-fold and 6.8-fold more respectively) (Fig. S6). Similarly, *trm8Δ* mutants expressing two copies of tP(AGG) had more tP(AGG) at 30°C and 38.5°C than the vector control, and the levels of the two control tRNAs were unchanged in all strains at both temperatures. We conclude that although levels of both tY(GUA) and tP(AGG) were reduced in *trm8Δ* mutants at elevated temperatures in both YES and EMMC media, tY(GUA) is the major physiologically important tRNA for these phenotypes.

Although tY(GUA) overexpression almost completely restored growth of *S. pombe trm8Δ* mutants in YES and EMMC media at 38°C and 39°C, expression of both tY(GUA) and tP(AGG) was required to completely suppress the growth defects in YES + glycerol media (Fig. S7). By contrast, overexpression of tY(GUA) and tP(AGG) had no effect on the known temperature sensitivity of *trm8Δ* mutants in YES media containing 5-FU (61, 62) (Fig. S7), perhaps due to reduced levels of Ψ and 5- methyluridine modifications.

### *dhp1* mutations suppress the *S. pombe trm8Δ* growth defect and restore tY(GUA) and tP(AGG) levels

To identify the mechanisms that restore growth to *S. pombe trm8Δ* mutants at elevated temperatures, we isolated and analyzed spontaneous suppressors of the temperature sensitivity. One major class of four *trm8Δ* suppressors were as temperature resistant as WT on YES and EMMC media, and nearly as resistant on YES + 5-FU media (Fig. 3A, S8). Genome sequencing revealed that these mutants each had distinct missense mutations in the *RAT1* ortholog *dhp1^+^*. The *dhp1* mutations each occurred in a highly conserved residue, based on an alignment of 18 *RAT1*/*dhp1^+^* eukaryotic homologs from multiple phyla (Fig. 3B), and are presumably partial loss of function mutations as *S. pombe dhp1^+^*, like *S. cerevisiae RAT1*, is an essential gene (63, 64).

**Fig. 3.**
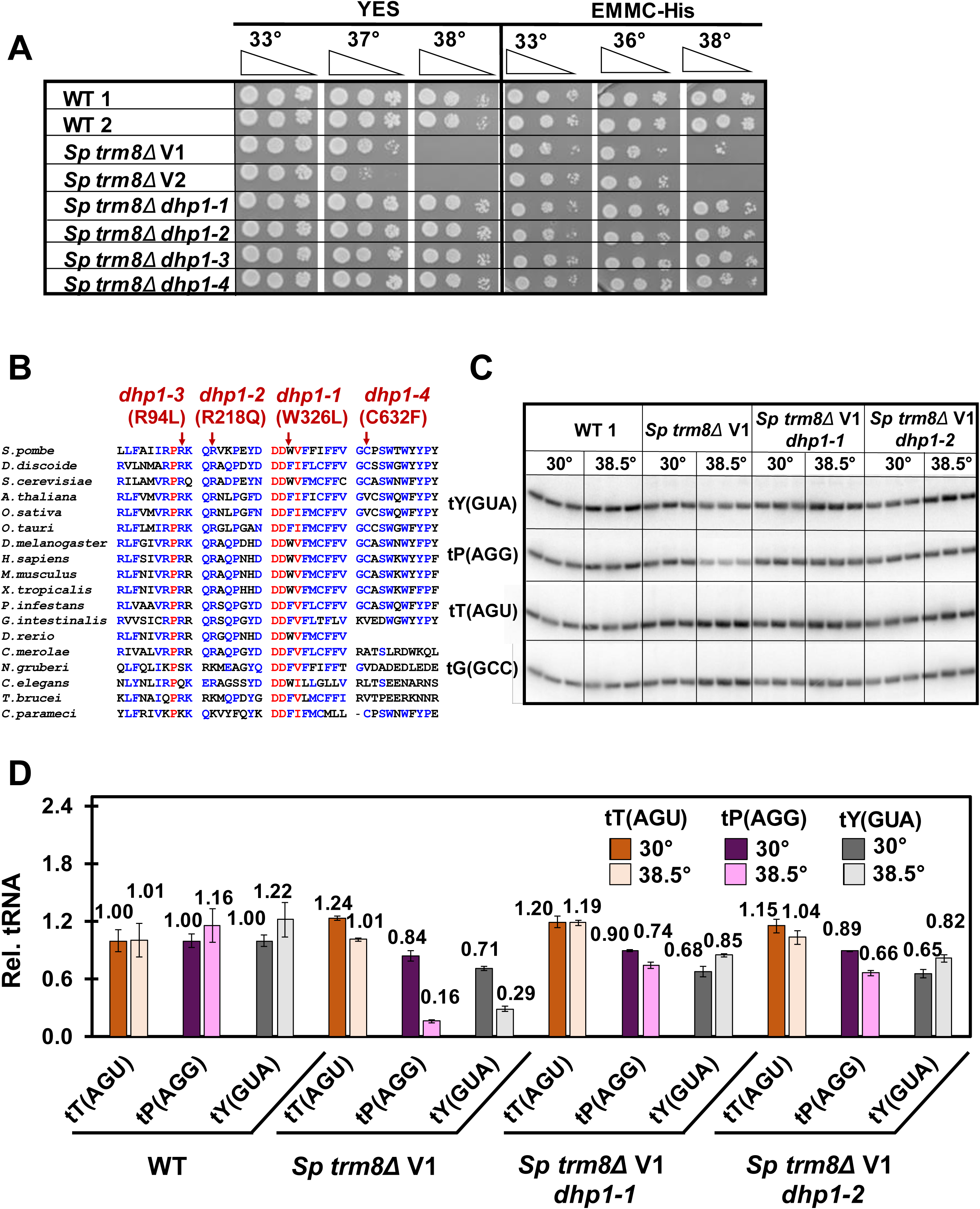
Mutations in *dhp1* suppress the temperature sensitivity of *S. pombe trm8Δ* mutants and restore tP(AGG) and tY(GUA) levels. (*A*) *dhp1* mutations restored growth of *trm8Δ* mutants at high temperature. Strains as indicated were grown overnight in YES media at 30°C and analyzed for growth as in Fig. 1B. (*B*) Mutations in *dhp1* that restored growth of *S. pombe trm8Δ* mutants reside in evolutionarily conserved residues. The amino acid sequence of *Sp* Dhp1 was aligned with putative Rat1/Dhp1 orthologs from 17 evolutionarily distant eukaryotes, using MultAlin (http://multalin.toulouse.inra.fr/multalin/) (133). red, > 90% conservation; blue, 50%-90% conservation. Alleles of *dhp1* mutations are indicated at the top. *(C*) Each of two *trm8Δ dhp1* mutants had restored tRNA levels in YES media at 38.5°C. Strains were grown in YES media at 30°C and shifted to 38.5°C for 8 hours, and RNA was isolated and analyzed by Northern blotting as in Fig. 2A. *(D)* Quantification of tRNA levels of *trm8Δ dhp1* mutants shown in Figure 3C. tRNA levels were quantified as in Fig. 2B. tT(AGU), brown; tP(AGG), purple; tY(GUA), gray; dark shades, 30°C; light shades, 38.5°C

Because we obtained four genetically independent *S. pombe trm8Δ dhp1* mutants and very few other mutations in the whole genome sequencing, it was highly likely that the *dhp1* mutations were responsible for the restoration of growth of *trm8Δ dhp1* mutants. Consistent with this, a plasmid expressing *dhp1*^+^ complemented the *S. pombe trm8Δ dhp1-1* suppressor, resulting in temperature sensitivity, but had no effect on WT or *trm8Δ* mutants (Fig. S9). Thus, we conclude that the *dhp1* mutations were responsible for the rescue of growth at high temperature.

As Dhp1 encodes a 5’-3’ exonuclease (65), it seemed highly likely that the *S. pombe trm8Δ dhp1* mutants prevented decay of tY(GUA) and tP(AGG) at non-permissive temperature. Indeed, we found that for each of two *dhp1* suppressors, tY(GUA) levels were almost completely restored at 38.5°C, from 29% in the *trm8Δ* mutant to 85% and 82% in the *trm8Δ dhp1-1* and *trm8Δ dhp1-2* strains respectively (Fig. 3C, 3D). Similarly, tP(AGG) levels were virtually completely restored at 38.5°C, from 16% in the *trm8Δ* mutant to 74% and 66% in the *trm8Δ dhp1* suppressors, and the levels of the control tRNA (tT(AGU)) was unaffected. Similar restoration of tY(GUA) and tP(AGG) levels was also observed in the two other *trm8Δ dhp1* suppressors at 38.5°C (Fig. S10). We conclude that, as for RTD in *S. cerevisiae* modification mutants (22, 39, 40), decay of tY(GUA) and tP(AGG) in *S. pombe trm8Δ* mutants occurs by 5’-3’ exonucleolytic degradation of tRNA, providing strong evidence for conservation of the RTD pathway in *S. pombe*.

### Mutations in the GAAC pathway suppress the *S. pombe trm8Δ* growth defect and restore tRNA levels

A second major group of six *S. pombe trm8Δ* suppressors was temperature resistant on YES and EMMC media, but sensitive on YES + 5-FU media, and genome sequencing showed that these suppressors each had distinct mutations in elements of the GAAC pathway (Fig. 4A, S11). Among these, we found three *trm8Δ* suppressors with *gcn2* mutations, one with a *gcn1* mutation, and two with *tif221* mutations, encoding the translation initiation factor eIF2Bα (Table S2). Each of these genes is known to be critical in the *S. cerevisiae* GAAC pathway (66–68), which is widely conserved in eukaryotes, including *S. pombe* and mammals (69–74). In this pathway, amino acid starvation leads to uncharged tRNAs that bind Gcn2 to activate its kinase domain, phosphorylation of eIF2α by Gcn2, and derepression of translation of the transcription factor Gcn4, resulting in increased transcription of nearly one tenth of the *S. cerevisiae* genes (66, 75–78). A similar massive transcription program change occurs in *S. pombe* after amino acid starvation (79).

**Fig. 4.**
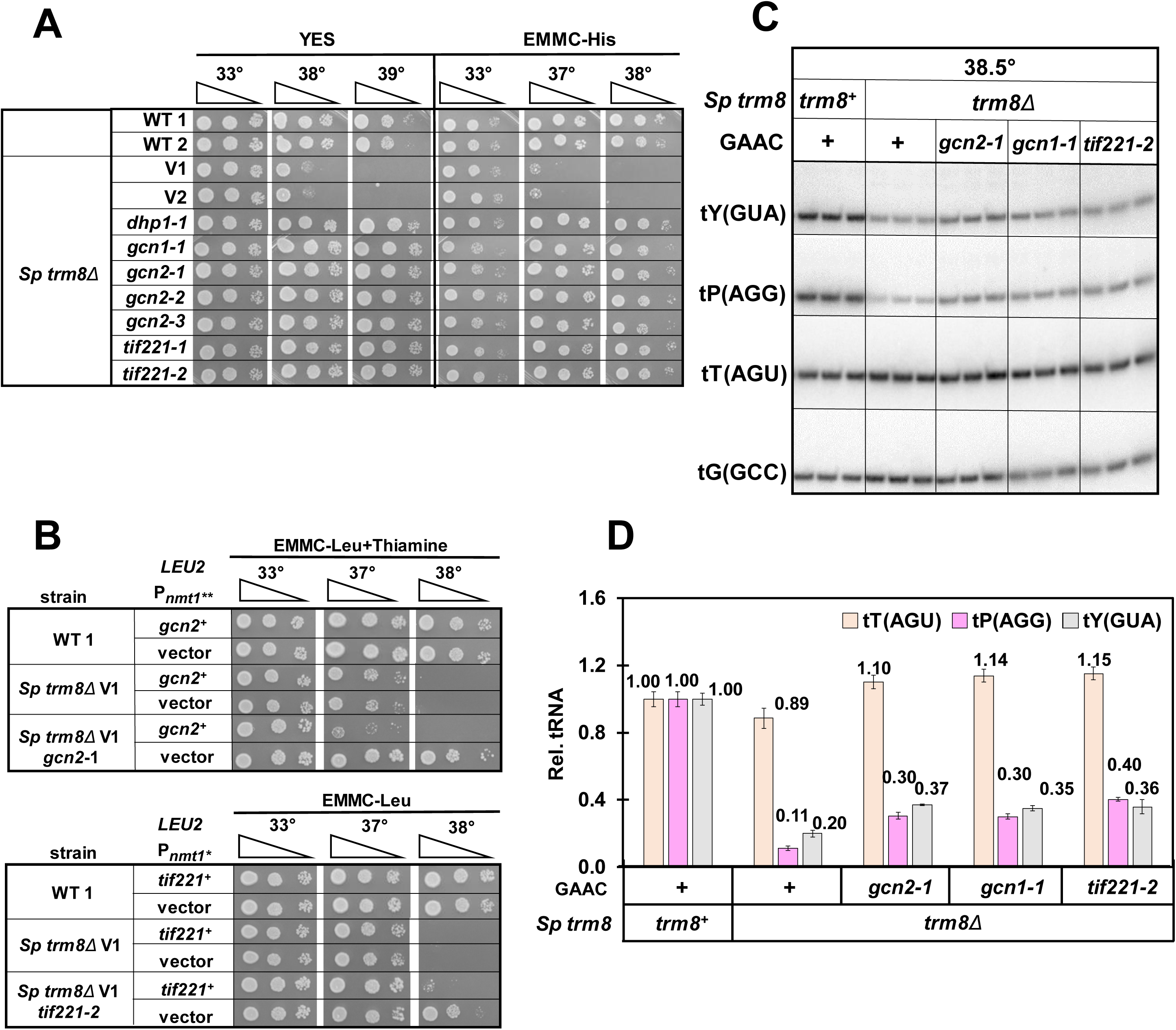
Mutations in the GAAC pathway suppress the temperature sensitivity of *S. pombe trm8Δ* mutants and restore tP(AGG) and tY(GUA) levels. (*A*) *gcn1, gcn2*, and *tif221* mutations each restored growth of *trm8Δ* mutants at high temperature. Strains as indicated were grown overnight in YES media at 30°C and analyzed for growth as in Fig. 1B. *(B*) Expression of *gcn2^+^* or *tif221^+^* complemented the suppression phenotype of *trm8Δ gcn2-1* or *trm8Δ tif221-2* mutants. *S. pombe trm8Δ gcn2-1* and *trm8Δ tif221-2* mutants expressing *gcn2^+^* and *tif221^+^* respectively, or a vector, were grown overnight in EMMC-Leu media at 30°C, and analyzed for growth as in Fig. 1B. Note that expression of *gcn2^+^* was kept to modest levels by adding thiamine to the media to partially suppress overexpression from the P*_nmt1_*\*\* promoter. *(C) gcn1, gcn2*, and *tif221* mutations each partially restored tY(GUA) and tP(AGG) levels of *trm8Δ* mutants. Strains were grown in YES media at 30°C and shifted to 38.5°C for 8 hours as described in Materials and Methods, and RNA was isolated and analyzed by northern blotting. *(D)* Quantification of tRNA levels of *trm8Δ* GAAC mutants shown in Figure 4C. tRNA levels were quantified as in Fig. 2B. tT(AGU), brown; tP(AGA), purple; tY(GUA), gray.

As expected, all *S. pombe trm8Δ* mutants with suppressing mutations in *gcn2*, *gcn1*, or *tif221*, grew poorly on media containing 3-Amino-1,2,4-triazole (3-AT) (Fig. S11), the classical inducer of the GAAC pathway (79–81). Furthermore, each of two *S. pombe trm8Δ* suppressors tested (with *gcn2-1* and *tif221-2* mutations) was complemented by re-introduction of the WT gene (Fig. 4B), and a re- constructed *trm8Δ gcn2Δ* strain was temperature resistant and sensitive to 3-AT (Fig. S12).

Consistent with their role as *S. pombe trm8Δ* suppressors, all six of the *trm8Δ* GAAC mutants had increased levels of tY(GUA) and tP(AGG) at high temperature. After growth in YES media at 38.5°C, all *trm8Δ* GAAC mutants showed a 1.7-fold to 2.1-fold increase in tY(GUA) levels compared to the parent *trm8Δ* mutant, and tP(AGG) levels were increased ∼ 3 fold, whereas the controls tT(AGU) and tV(AAC) did not have increased levels (Fig. 4C, 4D, S13). These results provided strong evidence that the effect of the GAAC mutants was to increase tRNA levels to restore growth.

### The temperature sensitivity of *S. pombe trm8Δ* mutants coincides precisely with the onset of GAAC activation and tY(GUA) decay

Since *S. pombe trm8Δ* suppressors in different components of the GAAC pathway all restored tY(GUA) and tP(AAG) levels, we speculated that *trm8Δ* mutants activated the GAAC pathway at non-permissive temperatures to somehow promote further loss of the tRNA. To establish the precise connection between growth, tRNA levels, and GAAC activation in *trm8Δ* mutants, we measured each parameter during liquid growth in rich media at a permissive temperature (30°C), and at three elevated temperatures: 36.5°C, 37.5°C, and 38.5°C. In this experiment, *trm8Δ* mutants grew virtually identically to WT control strains at 36.5°C and 37.5°C, and the growth defect was only obvious at 38.5°C (Fig. S14).

Strikingly, *S. pombe trm8Δ* mutants activated the GAAC pathway only at 38.5°C, the lowest temperature at which the growth defect was obvious. We measured GAAC activation by measuring mRNA levels of the known GAAC targets *lys4^+^* and *aro8^+^* (SPAC56E4.03) (79), which we had previously used (82). At 38.5°C in *trm8Δ* mutants, we observed a 7.1-fold increase in *lys4^+^* mRNA levels (relative to the standard *act1^+^*), compared to that from WT or *trm8Δ* mutants at 30°C (5.3 vs 0.74) (Fig. 5A). By contrast, we found no measurable change in *lys4^+^* mRNA levels in *trm8Δ* mutants grown at 36.5°C and 37.5°C, relative to that observed in *trm8Δ* mutants at 30°C, or in WT cells at any temperature. Moreover, the increase in relative *lys4^+^* mRNA levels in *trm8Δ* mutants in YES media at 38.5°C was almost as high as that observed in WT cells induced with 3-AT, and was completely eliminated in *trm8Δ gcn2-1* mutants. Examination of relative *aro8^+^* mRNA levels gave a similar result (Fig. S15): a substantial Gcn2-dependent increase in relative *aro8^+^* mRNA levels at 38.5°C in *trm8Δ* mutants, relative to 37.5°C (1.5 vs 0.45), and no change in relative *aro8^+^* mRNA levels at 37.5°C in *trm8Δ* mutants compared to WT (0.45 vs 0.44).

**Fig. 5.**
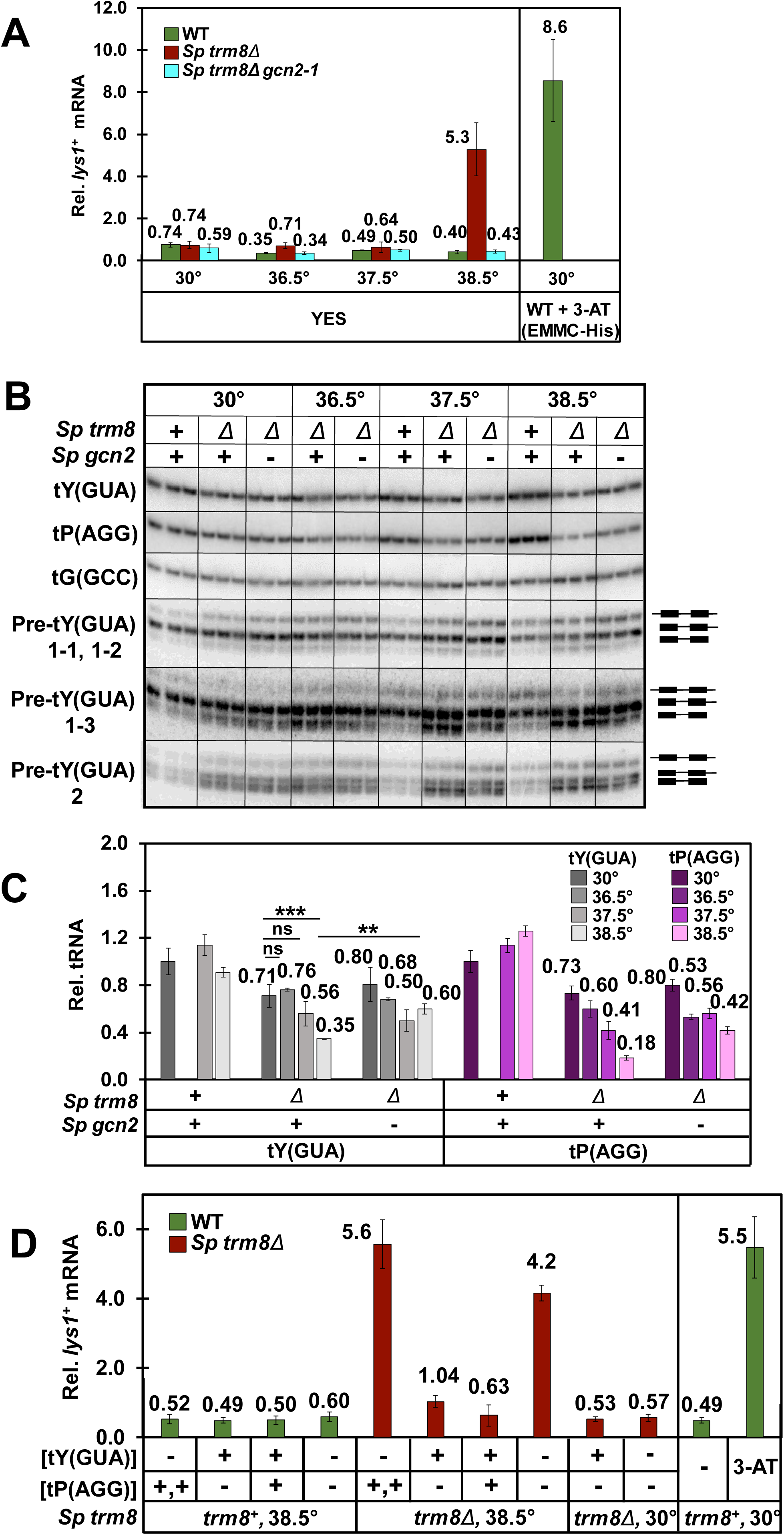
*S. pombe trm8Δ* mutant temperature sensitivity is associated with induction of the GAAC pathway due to Y(GUA) decay. *(A) trm8Δ* mutants induced *lys4^+^* mRNA expression at 38.5°C but not at 36.5°C or 37.5°C. Strains as indicated were grown in YES media at 30°C and shifted to 36.5°C, 37.5°C, or 38.5°C for 8 hours (Fig. S14), and bulk RNA was isolated and analyzed by RT-qPCR as described in Materials and Methods. The mRNA levels of *lys4^+^* were normalized to levels of *act1^+^*, a non-regulated control mRNA. WT, green; *Sp trm8Δ*, brown; *Sp trm8Δ gcn2-1,* light blue. Right side: GAAC induction of WT cells grown at 30°C in EMMC-His and treated with 10 mM 3-AT for 4 hours, evaluated in parallel. *(B)* A *trm8Δ gcn2-1* mutant had restored levels of tY(GUA) and tP(AGG) at 38.5°C. Bulk RNA from the growth done for Fig. 5A was used for the northern analysis. pre-tY(GUA) levels were analyzed using appropriate gene-specific probes (Table S5) for defined tY(GUA) species (59). Cartoons at the right indicate exons, heavy bars; 5’ leaders, 3’ trailers, and introns, light bars. *(C)* Quantification of tY(GUA) and tP(AGG) levels in WT, *trm8Δ* and *trm8Δ gcn2-1* mutants at different temperatures. tRNA levels were quantified as described in Fig. 2. *(D)* tY(GUA) overproduction repressed the GAAC induction of *trm8Δ* mutants at 38.5°C. Strains as indicated with plasmids expressing tY(GUA) and/or tP(AGG) were grown in EMMC-Leu media at 30°C and shifted to 38.5°C for 8 hours, and then RNA was isolated and *lys4^+^* mRNA levels were analyzed by RT-qPCR as described in Fig. 5A. Right side: GAAC induction of WT cells grown at 30°C in EMMC-His media, and treated with 10 mM of 3-AT for 4 hours, evaluated in parallel to other samples.

Consistent with the appearance of the *S. pombe trm8Δ* growth defect and the GAAC activation only at 38.5°C, tY(GUA) decay was only significant at 38.5°C (Fig. 5B, 5C), and the *gcn2-1* mutation significantly prevented that tY(GUA) decay. Moreover, the *gcn2-1* mutation appeared to exert its effect on tY(GUA) levels by preventing decay. Using intron probes for the four tY(GUA) genes, we found that levels of the primary transcript, corresponding to the largest species, were essentially unchanged in *trm8Δ gcn2-1* mutants, relative to *trm8Δ* mutants (Fig. 5B). As there was also no substantial reduction of any pre-tY(GUA) species in *trm8Δ* mutants at 38.5°C, relative to WT levels, we infer that tY(GUA) decay in *Sp trm8Δ* mutants occurred at the level of mature tRNA, as for RTD in *S. cerevisiae* (39). Indeed, the relative increase in several pre-tY(GUA) species in *trm8Δ* mutants at elevated temperatures, relative to WT, might imply a processing defect due to lack of m^7^G.

As tY(GUA) was the major physiologically relevant substrate in YES media (Fig. 2D), we speculated that at 38.5°C, tY(GUA) decay might be driving the GAAC activation associated with the *trm8Δ* growth defect. Alternatively, GAAC activation could be a consequence of both tY(GUA) and tP(AAG) decay, reduced tRNA charging associated with *trm8Δ* mutants at high temperature, or partly as a consequence of temperature stress itself, as a number of different stress conditions are known to activate the GAAC pathway (72, 74, 83, 84).

To determine the extent to which reduced tRNA levels activated the GAAC pathway, we examined GAAC induction of *S. pombe trm8Δ* strains after overproduction of tY(GUA) and tP(AGG), using the same samples we used to show that overproduction of tY(GUA) suppressed the *trm8Δ* temperature sensitivity (Fig. 2D, S6). As expected, relative *lys4^+^* mRNA levels were increased in *trm8Δ* [vector] strains grown at 38.5°C, compared to this strain at 30°C (4.17 vs 0.57, 7.3-fold) or to WT at 38.5°C (7.0 fold), indicating GAAC activation (Fig. 5D). Notably, *lys4^+^* mRNA levels were reduced 4.0-fold in the *trm8Δ* [tY(GUA)] strain compared to the corresponding *trm8Δ* [vector] strain(from 4.2 to 1.04), and were not reduced in the *trm8Δ* [tP(AGG)] strain. Based on these results, we conclude that reduced function of tY(GUA) is the primary cause of GAAC activation and temperature sensitivity of *trm8Δ* mutants at 38.5°C. As charging of tY(GUA) was distinctly but marginally reduced at 38.5°C relative to WT (Fig.S16), we cannot determine if the GAAC activation was due to reduced levels of tY(GUA), or to a combination of reduced levels and charging (69, 76, 80, 82, 85).

### In *S. cerevisiae*, mutation of the GAAC pathway exacerbates the effects of the RTD pathway

To investigate the evolutionary implications of the GAAC pathway on RTD, we examined the consequences of deletion of GAAC components in *S. cerevisiae trm8Δ trm4Δ* mutants, which are known to be temperature sensitive due to decay of tV(AAC) by the RTD pathway (18, 39). In contrast to our results in *S. pombe trm8Δ* GAAC mutants, deletion of *GCN1 or GCN2* exacerbated the temperature sensitivity of *S. cerevisiae trm8Δ trm4Δ* mutants in both rich (YPD) and minimal complete (SDC) media, and this phenotype was also observed upon deletion of the GAAC transcription factor *GCN4* (Fig. 6A). A similar exacerbated temperature sensitivity due to mutation of the GAAC pathway was also observed in other *S. cerevisiae* modification mutants known to be subject to RTD (22, 39), including *trm8Δ*, *trm1Δ*, *tan1Δ*, and *tan1Δ trm44Δ* mutants (Fig. S17).

**Fig. 6.**
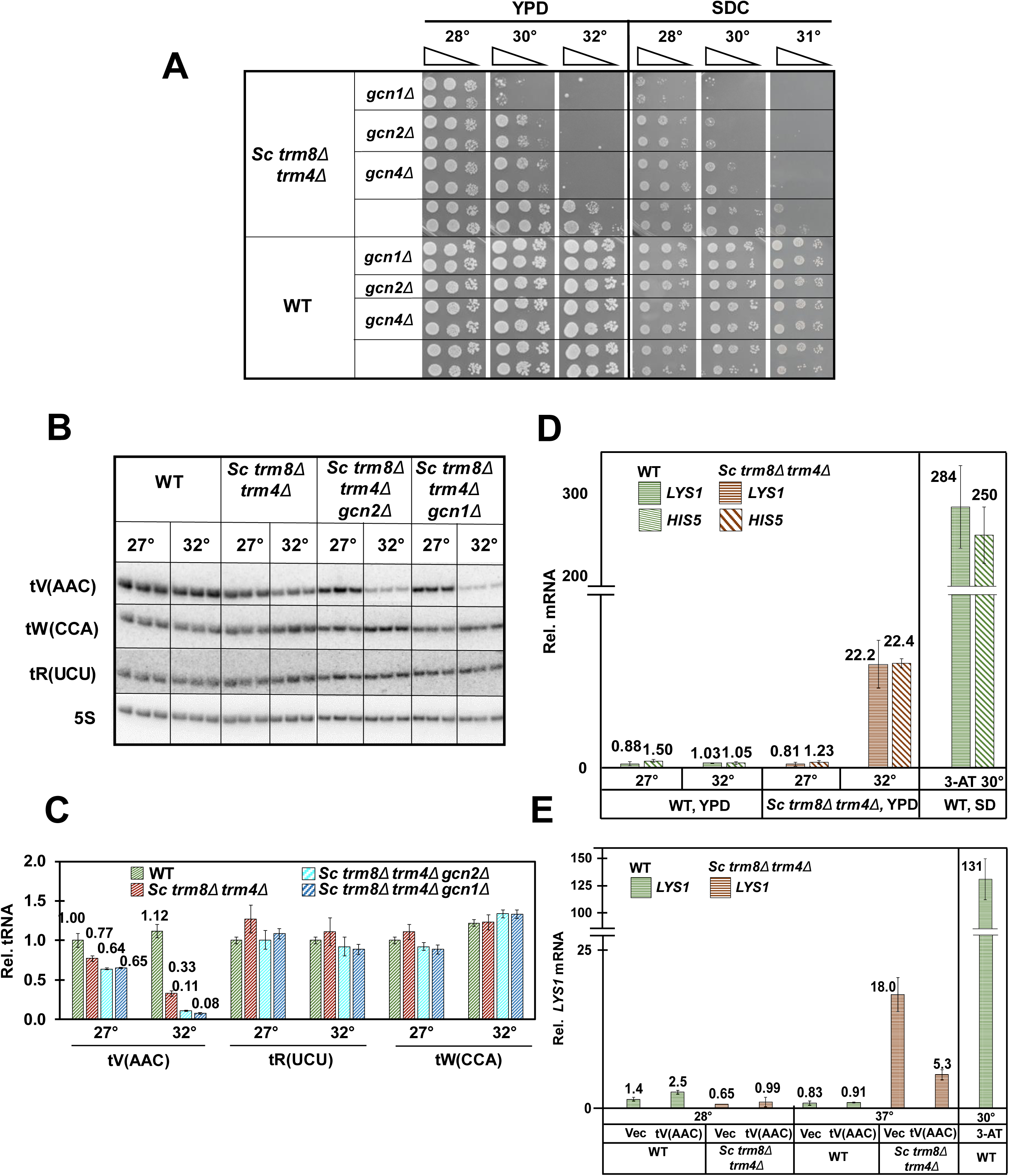
Mutations in the GAAC pathway exacerbate the temperature sensitivity of *S. cerevisiae trm8Δ trm4Δ* mutants as well as tV(ACC) decay. (*A*) Deletion of *GCN1, GCN2*, or *GCN4* exacerbated the temperature sensitivity of *trm8Δ trm4Δ* mutants in YPD and SDC media. Strains were grown overnight in YPD media at 27°C and analyzed for growth on YPD and SDC plates at different temperatures. *(B*) Deletion of *GCN1* or *GCN2* exacerbated tV(AAC) decay of *trm8Δ trm4Δ* mutants at 32°C. Strains were grown in YPD media 27°C, shifted to 32°C and harvested after 4 hours, and then bulk RNA was isolated and analyzed by Northern blotting. *(C)* Quantification of tRNA tV(AAC), tR(UCU) and tW(CCA) levels of the northern in Figure 6B. The bar chart depicts levels of tRNA species at 27°C or 32°C, relative to levels of that tRNA in the WT strain at 27°C (each value itself first normalized to levels of the control 5S rRNA). tRNA levels are indicated by diagonal hatching for WT (green); *S. cerevisiae* (*Sc*) *trm8Δ trm4Δ* (brown); *Sc trm8Δ trm4Δ gcn2Δ* (light blue); and *Sc trm8Δ trm4Δ gcn1Δ* (dark blue). *(D) trm8Δ trm4Δ* mutants induced the GAAC pathway at 32°C. Bulk RNA from the growth done for Fig. 6B was used for RT-qPCR analysis of levels of *LYS1* and *HIS5* mRNA, normalized to *ACT1*. mRNA levels are indicated by horizontal lines for *LYS1* and hatching for *HIS5*, WT (green); *Sc trm8Δ trm4Δ* (brown). Right side: Relative levels of *LYS1* mRNA of WT cells grown at 30°C in SD-His media, and then treated with 10 mM 3-AT for 1 hour, evaluated in parallel to other samples. *(E)* tV(AAC) overproduction repressed the GAAC induction of *trm8Δ trm4Δ* mutants at 36°C. Strains with plasmids as indicated were grown in SD-Ura media 27°C and shifted to 1 hour at 36°C and RNA was isolated and relative *LYS1* mRNA levels were analyzed by RT-qPCR as described in Fig. 6D. Right side: GAAC induction of WT cells grown and induced with 3-AT as described in Fig. 6D.

To determine if the exacerbated growth defect of *S. cerevisiae trm8Δ trm4Δ* GAAC mutants was due to exacerbated decay of tV(AAC), we analyzed tRNA levels after a four-hour temperature shift from permissive to non-permissive temperature (27°C to 32°C). Consistent with the exacerbated temperature sensitivity caused by the *gcn1Δ* and *gcn2Δ* mutations, tV(AAC) levels were further reduced in both *trm8Δ trm4Δ gcn1Δ* and *trm8Δ trm4Δ gcn2Δ* mutants at 32°C, compared to the *trm8Δ trm4Δ* mutant (11.8% and 18.5% vs 43.4%, relative to the values at 27°C) (Fig. 6B, 6C). As pre-tV(AAC) levels were unchanged at 32°C in the *trm8Δ trm4Δ gcn1Δ* mutants (Fig S18), we infer that tV(AAC) transcription was not affected by the *gcn1Δ* mutation, and that the enhanced loss of tV(AAC) was due to enhanced decay of mature tV(AAC). If so, then the enhanced temperature sensitivity of *trm8Δ trm4Δ* mutants due to a *gcn2Δ* or a *gcn4Δ* mutation was likely also due to enhanced decay of tV(AAC). Consistent with the conclusion that GAAC mutations enhanced RTD in *S. cerevisiae*, we found that a *met22Δ* mutation, known to prevent RTD, reversed the enhanced temperature sensitivity of a *trm8Δ trm4Δ gcn2Δ* strain relative to a *trm8Δ trm4Δ* mutant (Fig. S19).

### The temperature sensitivity and tV(AAC) decay of *S. cerevisiae trm8Δ trm4Δ* mutants coincides with GAAC activation

Since GAAC mutations exacerbated the RTD growth defect and enhanced the decay of an *S. cerevisiae trm8Δ trm4Δ* mutant at 32°C, it seemed likely that the GAAC pathway was activated in the *trm8Δ trm4Δ* mutant. To evaluate GAAC activation, we measured mRNA levels of the Gcn4 target genes *LYS1* and *HIS5* after the 4 hour temperature shift of *trm8Δ trm4Δ* mutants to 32°C, using the same RNA as in the Northern analysis of decay (Fig. 6B, 6C, S18). RT-qPCR analysis showed that *trm8Δ trm4Δ* mutants had a large increase in relative *LYS1* mRNA levels at 32°C, compared to 27°C (22.2 vs 0.81, 27.4-fold), or to WT cells at either 27°C or 32°C (Fig. 6D), showing that the GAAC activation was specific to the *trm8Δ trm4Δ* mutant at 32°C. We observed a similar activation of the *GCN4* target *HIS5* in the *trm8Δ trm4Δ* mutant at 32°C. This GAAC activation was correlated with a modest but distinct increase in uncharged tV(AAC) commensurate with the reduced tV(AAC) levels (Fig. S20). Furthermore, we found that overproduction of tV(AAC) in *trm8Δ trm4Δ* mutants suppressed induction of the GAAC pathway (Fig. 6E), showing that GAAC activation in *trm8Δ trm4Δ* mutants was due to the reduced function of tV(AAC). Thus, in both *S. cerevisiae trm8Δ trm4Δ* mutants and *S pombe trm8Δ* mutants, degradation of a single biologically relevant tRNA is the cause of GAAC induction, which then either promotes further decay in *S. pombe*, or prevents further decay in *S. cerevisiae* (Fig. 7).

**Fig. 7.**
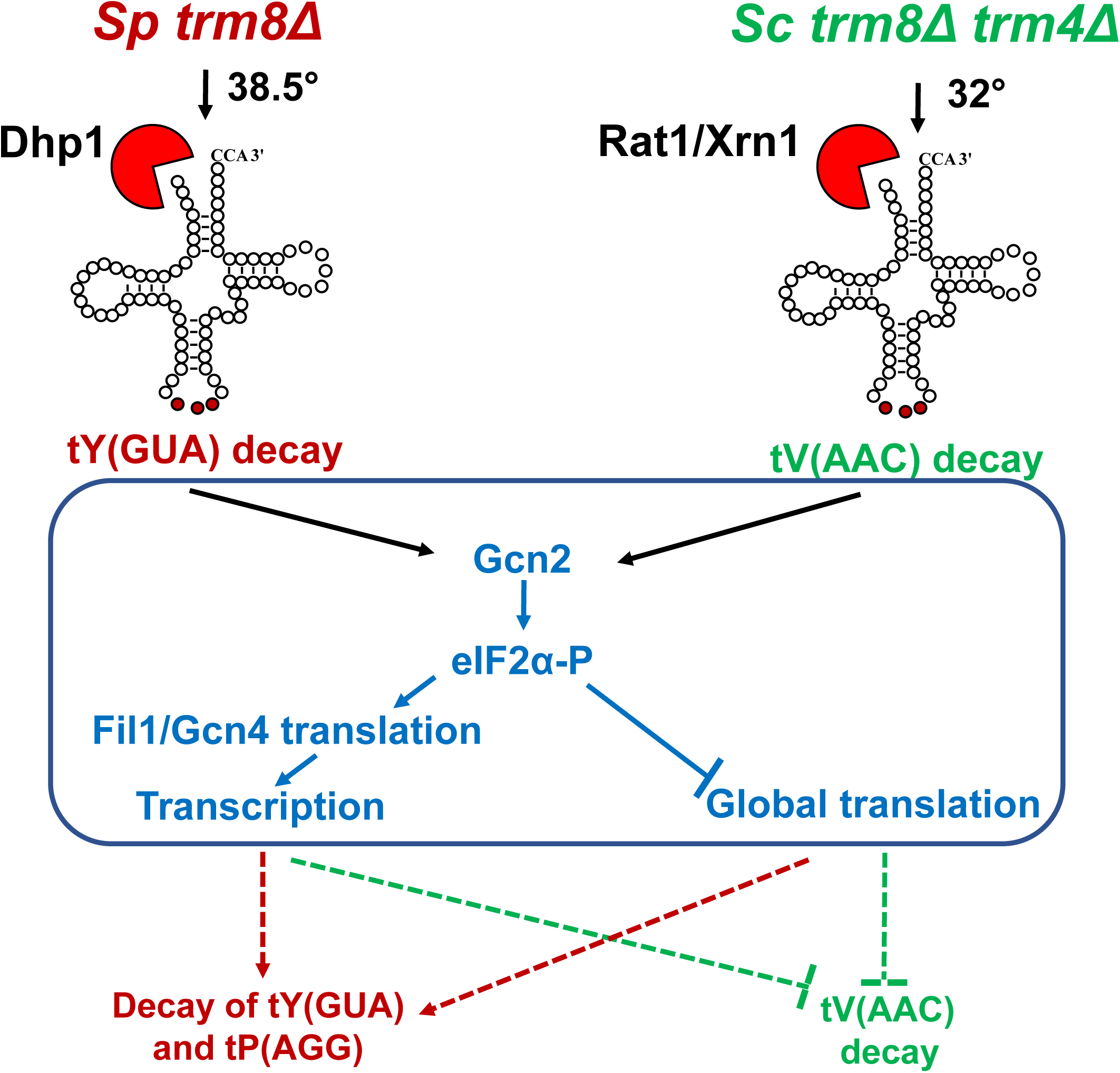
A model illustrating the interplay of the RTD pathway and GAAC induction in *S. pombe* and *S. cerevisiae*. Left: *S. pombe trm8Δ* mutants (red) trigger RTD of tY(GUA), leading to GAAC induction and further decay of tY(GUA) and tP(AGG). The increased tRNA decay resulting from GAAC induction could be due to the global reduction in translation and/or the transcription upregulation of Fil1 target genes. Right: *S. cerevisiae trm8Δ trm4Δ* mutants (green) trigger RTD of tV(AAC), leading to GAAC induction and inhibition of decay of tV(AAC). The reduced tRNA decay due to GAAC induction could be due to the reduced global translation and/or the transcription upregulation of Gcn4 target genes.

## Discussion

The results described here provide strong evidence that the RTD pathway is conserved between the distantly related species *S. cerevisiae* and *S. pombe*. We have shown that the temperature sensitivity of *S. pombe trm8Δ* mutants is due to reduced levels of tY(GUA) and to some extent tP(AGG), and is efficiently suppressed by mutations in the 5’-3’ exonuclease Dhp1 that concomitantly restore the levels of these tRNAs, strongly suggesting decay of tY(GUA) and tP(AGG) by the RTD pathway. As RTD is triggered in *S. cerevisiae* strains lacking m^2,2^G_26_ or ac^4^C_12_, as well as in strains lacking m^7^G_46_, (22, 39), we speculate that the RTD pathway will also act in *S. pombe* strains lacking other body modifications. Furthermore, given the large evolutionary distance between *S. cerevisiae* and *S. pombe,* we speculate that the RTD pathway is conserved throughout eukaryotes. The existence of a mammalian RTD pathway could explain the reduced levels of specific tRNA species in mouse strains lacking m^5^C in their tRNAs (33, 86) and in mouse embryonic stem cells lacking m^7^G in their tRNAs (46), and might explain other phenotypes associated with mutations in *METTL1* or *WDR4* (29, 30, 46, 87).

It is puzzling that although tP(AGG) is substantially more degraded than tY(GUA) in *S. pombe trm8Δ* mutants at elevated temperatures, the temperature sensitivity of the mutants in both rich and minimal media is primarily due to decay of tY(GUA). One possible explanation of this result is that tY(GUA) levels might be more limiting in the cell than tP(AGG) levels at elevated temperature, relative to the number of their respective cognate codons requiring decoding. This type of argument was advanced as a possible explanation for why the growth defects of i^6^A-lacking *S. pombe tit1Δ* mutants were rescued by increased expression of tY(GUA), but not by any of the other four Tit1 tRNA substrates (88). A second, and less likely, interpretation is that tP(AGG) decoding might be compensated by other tRNA^Pro^ isodecoders specific for the CCN codon box. This explanation is based on the finding that in *S. cerevisiae,* deletion of the two tP(AGG) genes is viable, implying that the 10 tP(UGG) isodecoders can decode all four proline CCN codons (89). However, as *S. pombe* has six tP(AGG) genes and only two tP(UGG) genes (as well as one tP(CGG)), it seems unlikely that the loss of almost all of the tP(AGG) in *trm8Δ* mutants can be efficiently compensated by the small number of tP(UGG) species (assuming that tRNA expression from each gene is comparable).

It is not immediately clear why tP(AGG) and tY(GUA) are the specific tRNAs subject to RTD in *S. pombe trm8Δ* mutants. Based on current understanding of RTD determinants in *S. cerevisiae*, RTD substrate specificity is determined by stability of the stacked acceptor and T-stem, with contributions to stability from the tertiary fold that are reduced in modification mutants, and some contributions from the other two stems, but not the anticodon loop (18, 22, 39–42). tP(AGG) may be an RTD substrate because it is predicted to have a less stable acceptor and T-stem than most Trm8 substrates (Table S1) (90). Furthermore, the destabilizing C_4_-A_69_ mismatch in the middle of the tP(AGG) acceptor stem might be expected to lead to increased local breathing at the 5’ end, which is likely important for recognition by the 5’-3’ exonucleases Xrn1 and Rat1, as the Xrn1 active site binds the three most 5’ nucleotides (91). However, it is more difficult to rationalize why tY(GUA) is a substrate for RTD in *trm8Δ* mutants, as its acceptor and T-stem are predicted to be moderately stable among Trm8 substrates. However, tY(GUA) does have a destabilizing N_27_-N_43_ pair (C_27_-U_43_ for 3 isodecoders, and U_27_-U_43_ for one isodecoder), which might reduce the stability of the tertiary fold by affecting stability of the adjacent tertiary 26-44 interaction (92–94).

It seems likely by analogy with RTD in *S. cerevisiae* that tRNA decay by Dhp1 in *S. pombe* involves retrograde transport of mature tRNA from the cytoplasm back to the nucleus. In *S. cerevisiae*, Rat1 is nuclear (95) and catalyzes a substantial amount of the decay of mature tV(AAC) in *S. cerevisiae trm8Δ trm4Δ* mutants (39), implying that the retrograde transport pathway is required to deliver the tV(AAC) substrate to the nucleus (5, 96–99). We argued above that mature tY(GUA) was the actual substrate for decay in *S. pombe trm8Δ* mutants. As pre-tRNA splicing in *S. pombe* is known to initiate on the surface of mitochondria (100), mature tY(GUA) is almost certainly transported back to the nucleus by the tRNA retrograde pathway to be degraded by Dhp1(64, 101). Retrograde transport could occur in *S. pombe trm8Δ* mutants driven in part by the hypomodification, as speculated previously in *S. cerevisiae* and shown for *S. cerevisiae trm1Δ* mutants (99).

It seems likely that the 5-FU sensitivity of *S. pombe trm8Δ* mutants is due to decay of multiple tRNA species in the presence of the drug, caused by the reduced levels of Ψ and m^5^U (48–50), in addition to the lack of m^7^G. This interpretation is consistent with the lack of suppression of the 5-FU sensitivity of *trm8Δ* mutants by tY(GUA) and tP(AGG), and its almost complete suppression by *dhp1* mutations, and is consistent with the enhanced 5-FU sensitivity of a number of tRNA body modification mutants in *S. cerevisiae* (61).

Our finding that loss of function of tRNA due to tRNA decay is itself the trigger for induction of the GAAC response in both *S. pombe trm8Δ* mutants and *S. cerevisiae trm8Δ trm4Δ* mutants suggests an intimate relationship between reduced tRNA function and GAAC activation. The loss of functional tRNA is the proximal cause of GAAC induction, because in each organism the GAAC pathway is activated at the lowest temperature at which tRNA decay and a growth defect is observed, and in each organism overproduction of the physiologically relevant tRNA represses GAAC induction. The GAAC pathway has previously been implicated in the biology of a number of anticodon loop modifications (82, 102, 103). Robust constitutive GAAC induction is observed in *S. cerevisiae* and *S. pombe trm7Δ* mutants (lacking Nm_32_ and Nm_34_) and *S. cerevisiae pus3Δ* mutants (lacking Ψ_38_ and Ψ_39_) (82, 103), each of which has a constitutive growth defect (104, 105), and GAAC induction is known to be Gcn2-dependent in *S. cerevisiae trm7Δ* mutants (82). By contrast, *S. cerevisiae* mutants lacking either the mcm^5^U or the s^2^U moiety of mcm^5^s^2^U induce the GAAC pathway independently of Gcn2 at 30°C (102) and are temperature sensitive at 37°C (106). Our finding that *S. pombe trm8Δ* and *S. cerevisiae trm8Δ trm4Δ* mutants each trigger Gcn2-dependent GAAC induction only at the temperature that the growth defect is observed is consistent with Gcn2-dependent GAAC induction in *S. cerevisiae* anticodon loop modification mutants with a constitutive growth defect.

It is striking that the induction of the GAAC response due to tRNA decay in *S. pombe trm8Δ* mutants and *S. cerevisiae trm8Δ trm4Δ* mutants has opposite consequences in each organism. In *S. pombe trm8Δ* mutants, activation of the GAAC response exacerbates the growth defect, as mutation of any of three components (*gcn1*, *gcn2*, or *tif221*) protects against decay. Activation of the GAAC pathway is also part of the reason that *S. cerevisiae trm7Δ* mutants grow poorly (82), and defects in the integrated stress response pathway (ISR) in humans are implicated in disease phenotypes (107–109). By contrast, in *S. cerevisiae trm8Δ trm4Δ* mutants, activation of the GAAC response rescues the growth defect, as deletion of any of three GAAC components *(gcn1Δ, gcn2Δ*, or *gcn4Δ*) exacerbates the growth defect. Although this rescue of RTD in *S. cerevisiae* by GAAC activation is opposite to that in *S. pombe*, it is consistent with a concerted stress response. Moreover the finding that *S. cerevisiae gcn4Δ* mutations exacerbated RTD nearly as much as *gcn1Δ* or *gcn2Δ* mutations has mechanistic implications. Deletion of *GCN1* or *GCN2* each prevent sensing of tRNA status, and the consequent eIF2α phosphorylation, reduced translation initiation, and reduced global translation. However, as *GCN4* is downstream of the sensing machinery, we infer that the *gcn4Δ*-mediated GAAC effects on RTD in *S. cerevisiae* are mostly due to lack of transcription activation by Gcn4, rather than the increased eIF2α phosphorylation leading to the reduction in translation initiation and overall translation.

Our evidence that GAAC activation has opposite consequences in *S. pombe* and *S. cerevisiae* is likely due to the differential GAAC response in genes known to regulate RTD. Although in each organism the GAAC pathway is known to regulate the transcription of more than 500 genes (75, 77, 79, 110), there are distinct differences in the GAAC response in the two species. For example, it is known that the GAAC response to amino acid starvation results in repression of methionine synthesis genes in *S. pombe* but induction of these genes in *S. cerevisiae* (79). As a *met22Δ* mutation is known to inhibit RTD in *S. cerevisiae*, this opposite GAAC activation effect on methionine genes in the two organisms is in the wrong direction to explain the opposite RTD effects of the GAAC pathway. Other possible explanations for the differential effects of the GAAC pathway on RTD include differential regulation of the synthesis or biochemical activity of RTD regulators such as EF1A, aminoacyl tRNA synthetases, pol III transcription (22, 111), or 5’-3’ exonucleases (39, 42), as well as changes in any number of indirect effectors affecting overall levels or availability of tRNA and/or nucleases. Moreover, the overall stress response pathways are substantially different between *S. cerevisiae* and *S. pombe*. In *S. cerevisiae,* Gcn2 is the sole eIF2a kinase regulating stress responses (78, 112, 113), whereas in *S. pombe* three different eIF2a kinases (Gcn2, Hri1 and Hri2) (114) each respond to a diverse set of stress treatments (72, 74, 83, 84, 107). The differences in kinases affecting eIF2α phosphorylation implies substantial differences between the two species in regulation of all sorts of combinations of stress response, which might be occurring at elevated temperature when tRNA decay is occurring (115).

The results outlined here underscore that GAAC activation occurs in *S. cerevisiae* and *S. pombe trm8Δ* modification mutants precisely at the point of observed growth stress due to tRNA decay, albeit with different effects in *S. pombe trm8Δ* mutants and *S. cerevisiae trm8Δ trm4Δ* mutants. These results, coupled with the constitutive GAAC activation in *S. pombe* and *S. cerevisiae trm7Δ* mutants and *S. cerevisiae pus3Δ* mutants (82, 103), fuel speculation that the GAAC response will also be activated in mammals and other eukaryotes with tRNA modification mutants or mutations that result in growth phenotypes due to reduced tRNA function.

As Trm8 is phosphorylated and likely inactivated by treatment of HEK293 cells with insulin-like growth factor-1 (116), it seems plausible that if RTD is conserved in mammals, such m^7^G modification dynamics could be used to regulate tRNA levels physiologically. There is growing evidence that levels of a number of modifications are under dynamic control in different conditions (117, 118). There is also evidence that regulation of tRNA expression plays an important role in differentiation and proliferation, and is also a characteristic of breast cancer (119–123). For example, tRNA^Arg^ iso-acceptors have different expression in differentiated vs proliferating cells (122) and tR(UCU) iso-decoders show tissue specific regulation of expression (124). It remains to be determined if regulated changes in expression, phosphorylation, or biochemical activity of METTL1 or WDR4 result in altered m^7^G levels and consequent changes in tRNA levels that physiologically regulate expression.

## Materials and Methods

### Yeast Strains

*S. pombe* haploid WT and two independent *S. pombe trm8Δ::kanMX* strains were derived from SP286 (*ade6-M210/ade6-M216, leu1-32/leu1-32, ura4-D18/ura4-D18 h+/h+*)(125), and were obtained from Dr. Jeffrey Pleiss. *S. pombe trm8Δ::kanMX* strains were also generated from haploid WT strains by PCR amplification of *trm8Δ::kanMX* DNA, followed by linear transformation using lithium acetate (126). *S. cerevisiae* deletion strains are shown in Table S3, and were constructed by linear transformation with PCR amplified DNA from the appropriate knockout strain (127). All strains were confirmed before use by PCR amplification.

### Plasmids

Plasmids used in this study are listed in Table S4. AB553-1 was constructed by insertion of a NotI restriction site between the P*_nmt1_* promoter and the PstI site of the pREP3X plasmid. The *S. pombe* plasmid expressing *S. pombe* P*_trm8+_ trm8^+^* (ETD 67-1) was constructed by inserting PCR amplified DNA genomic DNA (including 1000 bp upstream and 1000 bp downstream) into the NotI and XhoI sites of AB 553-1, removing the P*_nmt1_* promoter. Plasmids expressing *S. pombe* P*_dhp1_ dhp1^+^* or *S. pombe* tRNA genes were constructed using the same approach, including ∼ 300 bp upstream and 300 bp downstream for the tRNA genes. *S. pombe* plasmids expressing P*_nmt1**_ gcn2^+^* (low strength, no message in thiamine) or P*_nmt1*_ tif221^+^* (media strength, no message in thiamine) were constructed by PCR amplification of the respective coding sequence from *S. pombe* WT genomic DNA (including introns), and insertion into the XhoI and BamHI sites of the pREP81X or pRE41X vectors respectively.

### Yeast media and growth conditions

*S. pombe* strains were grown at desired temperatures in rich (YES) media (containing 0.5% yeast extract, 3% glucose, and supplements of 225 mg/l of adenine, uracil, leucine, histidine and lysine), or Edinburgh minimal media (EMM) containing glucose and the same supplements, as well as similar amounts of relevant auxotrophic requirements. Minimal complete (EMM-C) media was supplemented with 225 mg/l of all amino acids, adenine, and uracil, as well as 100 mg/l of para-amino benzoic acid and inositol, and 1125 mg/l of leucine for Leu^-^ auxotrophs (79). For temperature shift experiments, cells were grown in YES or EMMC media at 30°C to OD_600_ ∼ 0.5, diluted to ∼ 0.1 OD in pre-warmed media at the desired temperature, grown to OD ∼ 0.5, harvested at 4°C, washed with ice cold water, frozen on dry ice, and stored at −80°C. To select spontaneous suppressors of *S. pombe trm8Δ* mutants, cells were grown overnight in YES media at 30°C and ∼10^7^ cells were plated on YES media plates at 38°C and 39°C. *S. cerevisiae* strains were grown in rich (YPD) media (containing 1% yeast extract, 2% peptone, 2% dextrose, and 80 mg/L adenine hemisulfate), or minimal complete (SDC) media (128) as indicated, and temperature shift experiments were performed as described for *S. pombe*. All experiments with measurements were performed in biological triplicate, unless otherwise noted.

### Bulk RNA preparation and northern blot analysis

For northern analysis, 2 or 3 biological replicates were grown in parallel, and then bulk RNA was isolated from ∼ 3-5 OD pellets using glass beads and phenol (129) (for *S. pombe*) or hot phenol (for *S. cerevisiae*), resolved on a 10% polyacrylamide (19:1), 7M urea, 1X TBE gel, transferred to Amersham Hybond-N^+^ membrane, and analyzed by hybridization to 5’ ^32^P-labeled DNA probes (Table S5) as described (18). For analyzing tRNA charging levels of both *S. pombe* and *S. cerevisiae*, RNA was prepared under acidic conditions (pH 4.5), resolved on a 6.5 % polyacrylamide (19:1), 8 M urea, 0.1 M sodium acetate (pH 5.0) gel at 4°C, and analyzed as described. (18).

### Quantitative RT-PCR analysis

Strains were grown in triplicate to log phase and bulk RNA was prepared from 2-5 OD pellets using acid washed glass beads and phenol. Then, RNA was treated with RQ1 RNase-free DNase (Promega), reverse transcribed with Superscript II Reverse Transcriptase, and quantitative PCR was performed on the cDNA as previously described (130).

### Isolation and purification of tRNA

*S. pombe* WT and *trm8Δ* mutant strains were grown to ∼ 0.5 OD in YES media at 30°C. Then bulk low molecular weight RNA was extracted from ∼ 300 OD of pellets by using hot phenol, and tRNAs were purified using 5’-biotinylated oligonucleotides complementary to the corresponding tRNAs (Table S6) as previously described (131).

### HPLC analysis of nucleosides of purified tRNA

Purified tRNAs (∼ 1.25 µg) were digested to nucleosides by treatment with P1 nuclease, followed by phosphotase, as previously described (131), and nucleosides were analyzed by HPLC at pH 7.0 as previously described (132).

### Whole genome sequencing

Whole genome sequencing was performed by the University of Rochester Genomics Center at a read depth of 20-110 per genome nucleotide.

## Acknowledgements

We are grateful to Dr. Jeffrey Pleiss for the gift of *S. pombe* strains and invaluable help in preparing samples for whole genome sequencing. We also thank Dr. Elizabeth Grayhack, and the members of the Phizicky lab and Grayhack lab for valuable discussions and comments during the course of this work. This research was supported by NIH Grant GM052347 to E.M.P.

## Supplementary Figure Legends

**Fig. S1.**
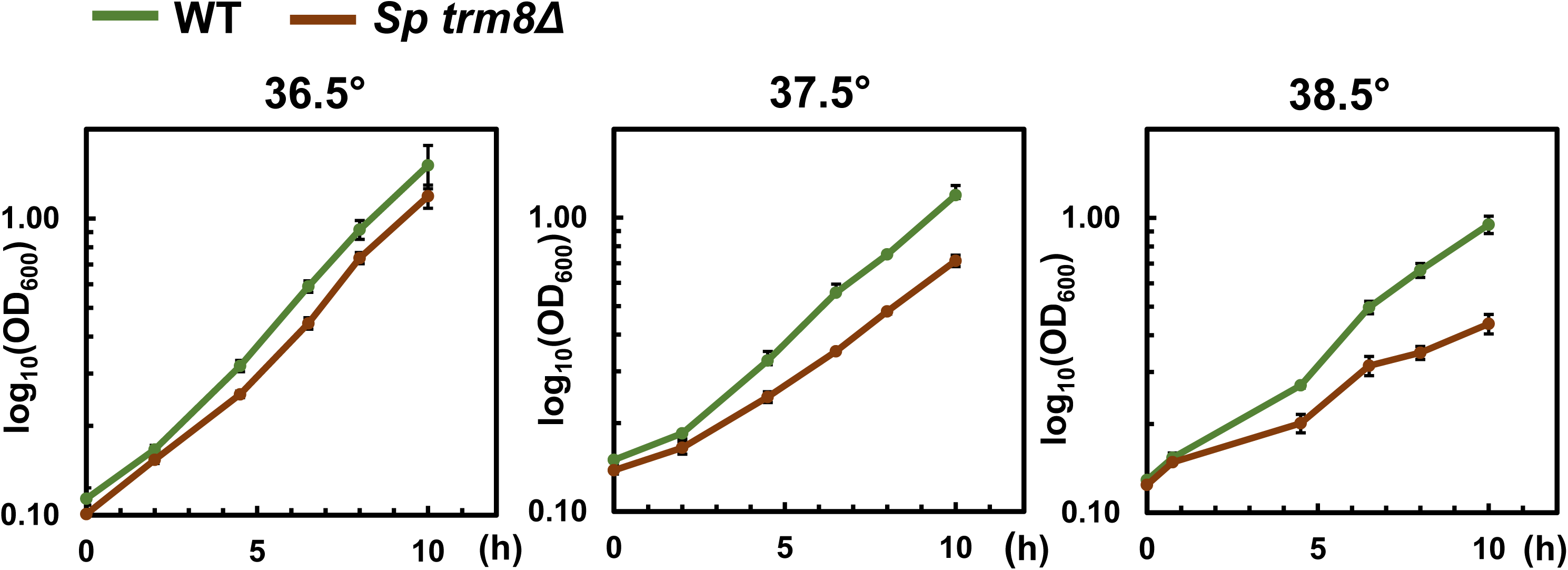
*S. pombe trm8Δ* mutants showed a temperature sensitive growth defect in liquid YES media. Strains were grown in YES media at 30°C, shifted to the indicated temperatures, and then growth was monitored for 8 hours before harvest as described in Materials and Methods, and tRNA analysis as done in Fig. 2A, 2B. WT, green; *Sp trm8Δ,* brown.

**Fig. S2.**
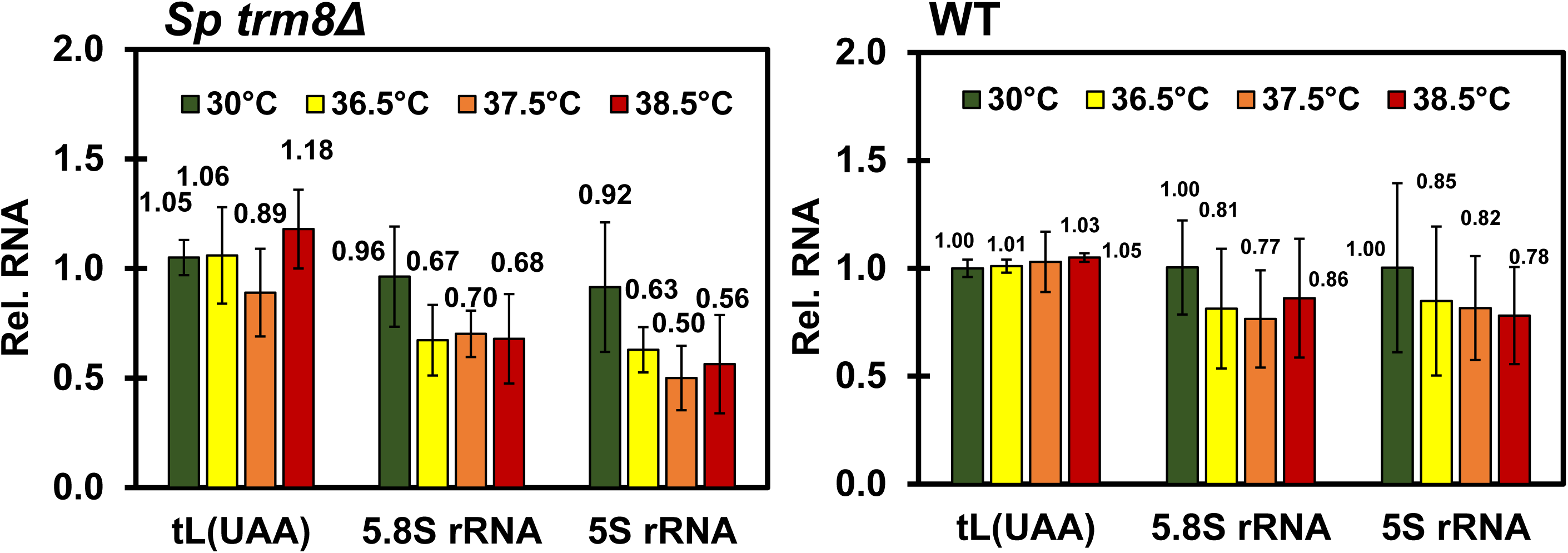
*S. pombe trm8Δ* mutants had reduced 5S rRNA and 5.8S rRNA levels at higher temperatures. The northern blot shown in Fig. 2A was used to analyze the non-Trm8 substrate tL(UAA), 5S rRNA, and 5.8S rRNA. The bar chart depicts levels of RNA species at each temperature, relative to their levels in the WT strain at 30°C (each value itself first normalized to levels of the control non-Trm8 substrate tG(GCC)). 30°C, green; 36.5°C, yellow; 37.5°C, orange; 38.5°C. red.

**Fig. S3.**
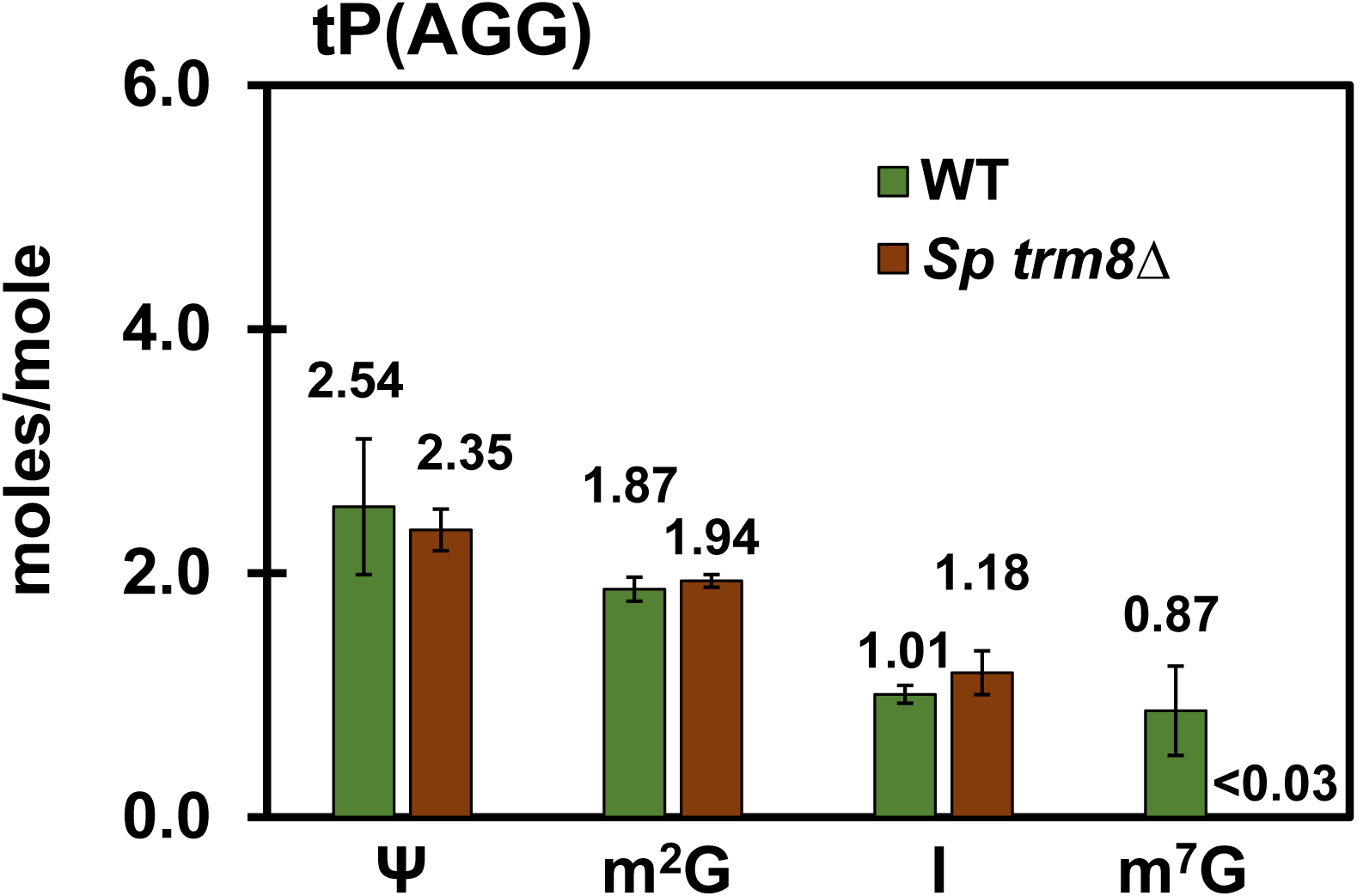
tP(AGG) of *S. pombe trm8Δ* mutants had no detectable m^7^G. *trm8Δ* mutants and WT cells were grown in YES media at 30°C and tP(AGG) was purified and analyzed for modifications as in Fig. 1A.

**Fig. S4.**
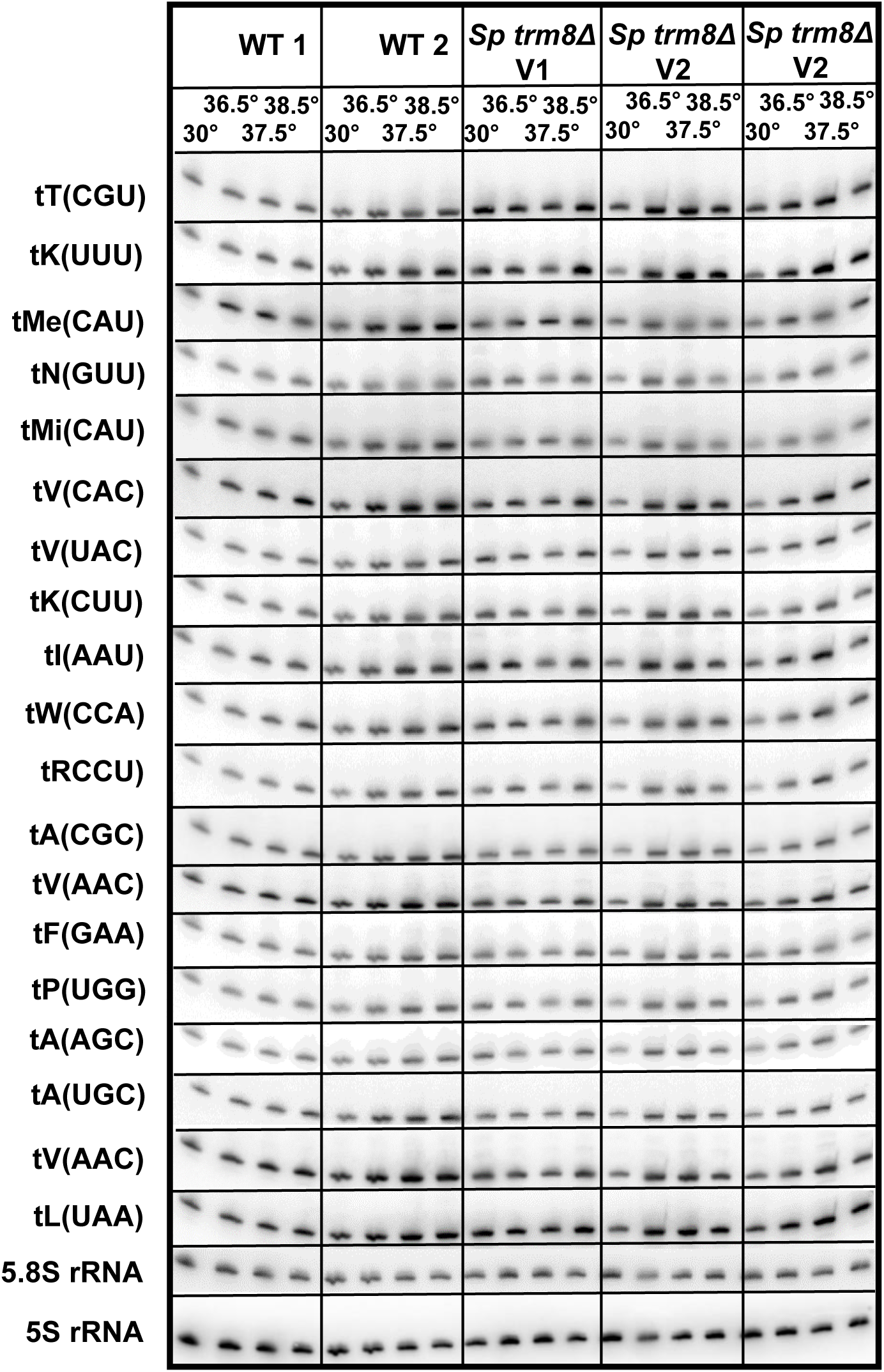
Northern analysis of all Trm8 substrates (except tP(AGG), tY(GUA), and tT(AGU)) in *S. pombe trm8Δ* and WT cells after shift from 30°C to 36.5°C, 37.5°C, and 38.5°C. The northern blot shown in Fig. 2A was continued to analyze levels of all other predicted Trm8 substrate tRNAs, as well as the non-Trm8 substrate tL(UAA), 5S RNA, and 5.8S RNA, in WT and *trm8Δ* mutants at different temperatures.

**Fig. S5.**
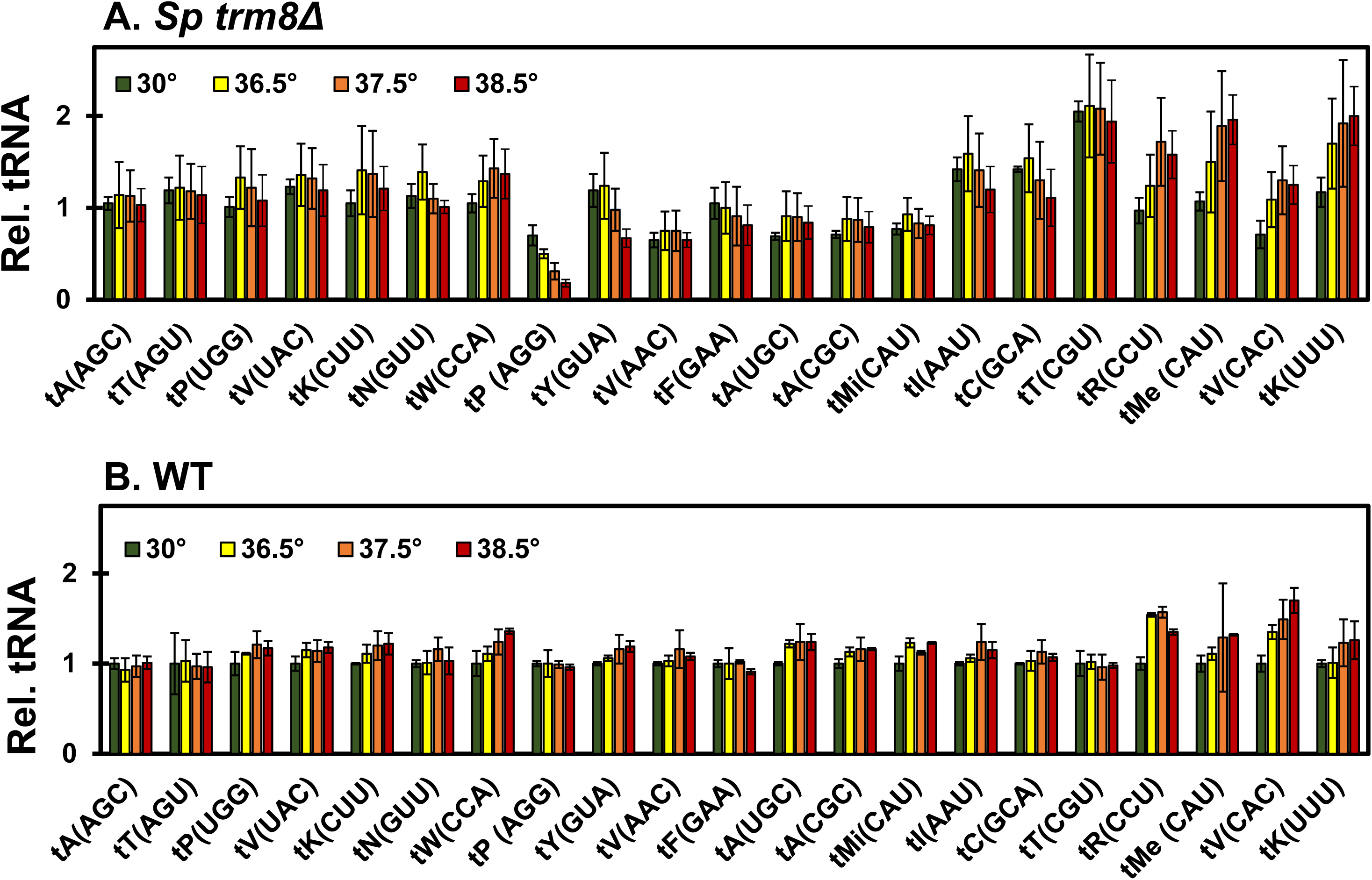
Among 21 predicted Trm8 substrate tRNAs, only tP(AGG) and tY(GUA) had reduced levels in *S. pombe trm8Δ* mutants at elevated temperatures. tRNA levels were quantified, relative to tG(GCC), in *trm8Δ* mutants and WT cells at different temperatures, for all predicted Trm8 substrates, as described in Fig. 2B. Note that data from Fig. 2B is also included here for completeness.

**Fig. S6.**
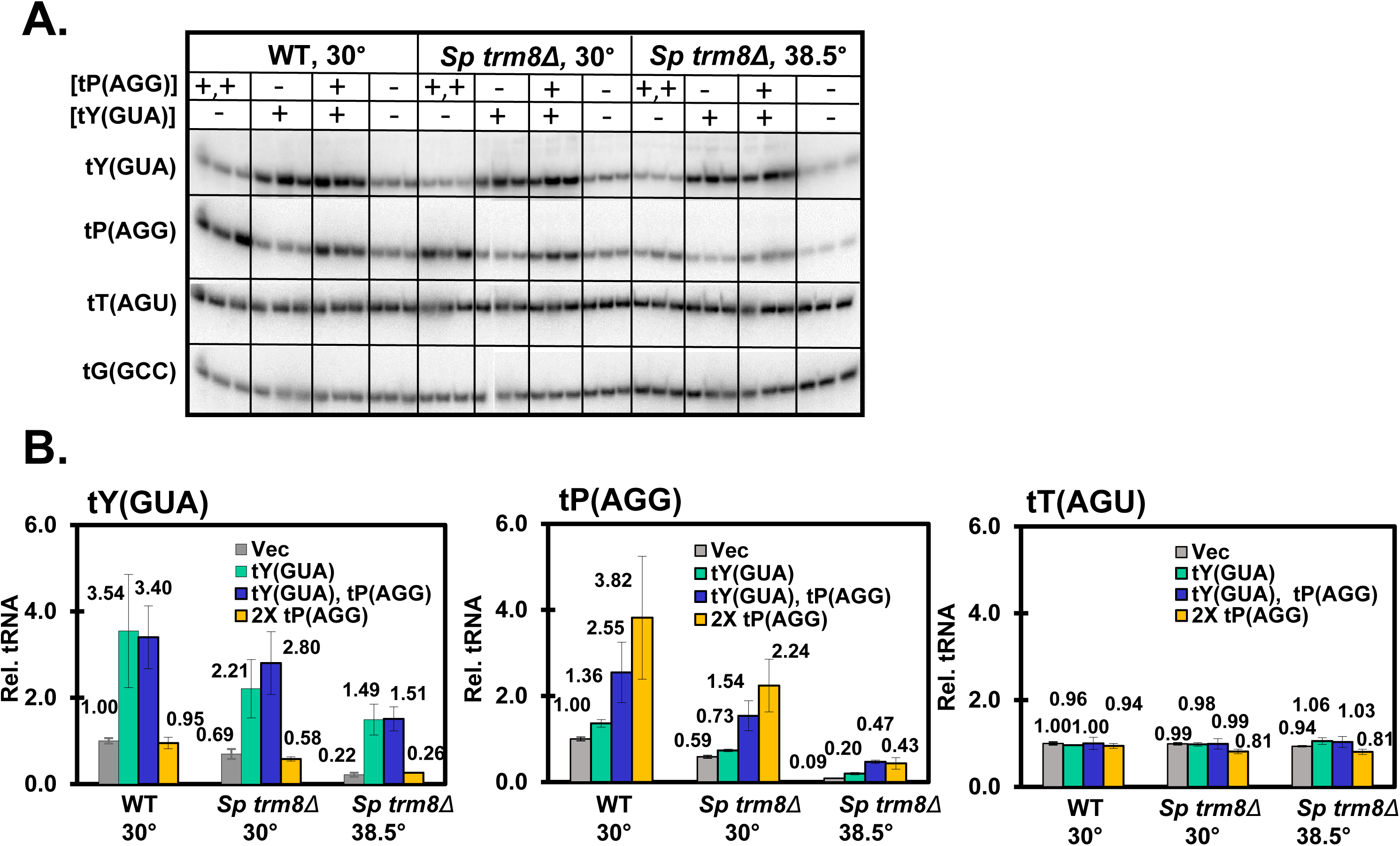
A. Overproduction of tY(GUA) and tP(AGG) resulted in increased levels of the corresponding tRNAs in *S. pombe trm8Δ* mutants and WT cells. Strains with plasmids as indicated were grown in EMMC-Leu media at 30°C and shifted to 38.5°C for 8 hours, and then RNA was isolated and analyzed by Northern blotting as in Fig. 2A. B. Quantification of tRNA levels in *S. pombe trm8Δ* mutants and WT cells overproducing tY(GUA) or tP(AGG). Quantification was done as in Fig. 2B.

**Fig. S7.**
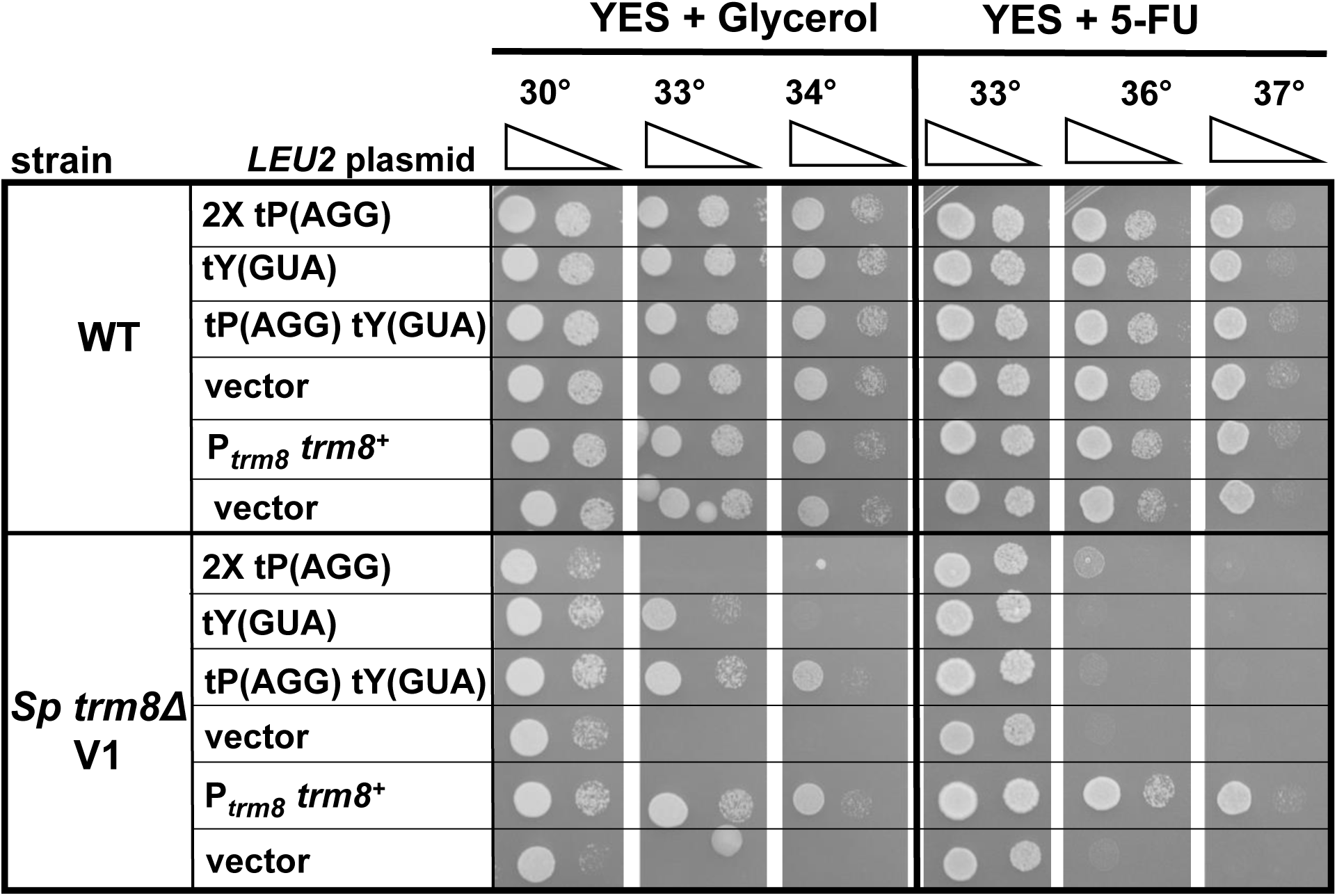
Overproduction of both tY(GUA) and tP(AGG) fully restored growth of *S. pombe trm8Δ* mutants in YES + glycerol media, but not in YES media containing 5-FU. Strains grown for Fig. 2D were analyzed for growth on plates containing YES media with 3% glycerol (instead of 3% glucose) and YES media with 5-FU (30 µg/ml).

**Fig. S8.**
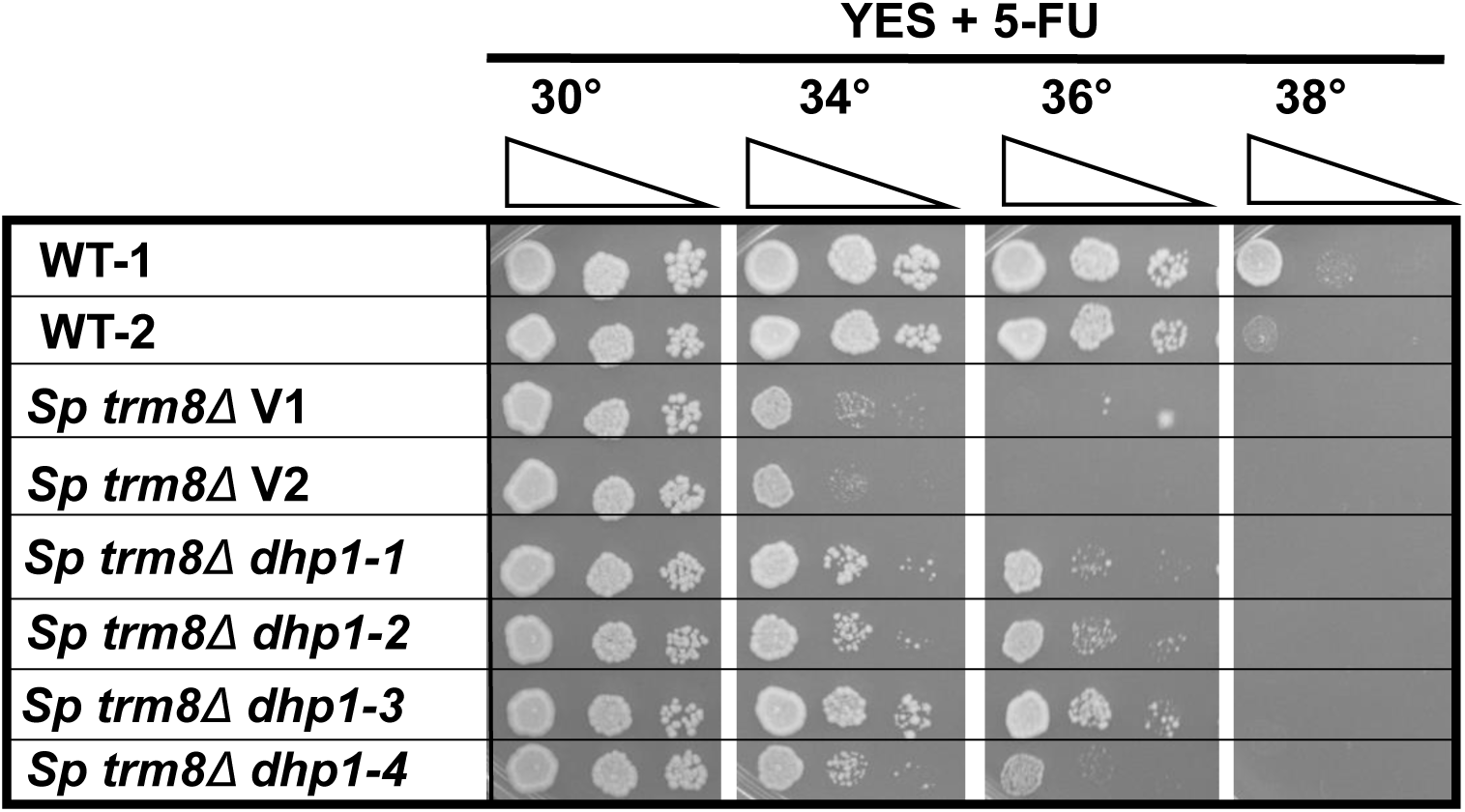
*dhp1* mutations restored growth of *S. pombe trm8Δ* mutants in YES + 5-FU media. Strains grown for Fig. 3A were analyzed for growth on YES + 5-FU (30 µg/ml) plates.

**Fig. S9.**
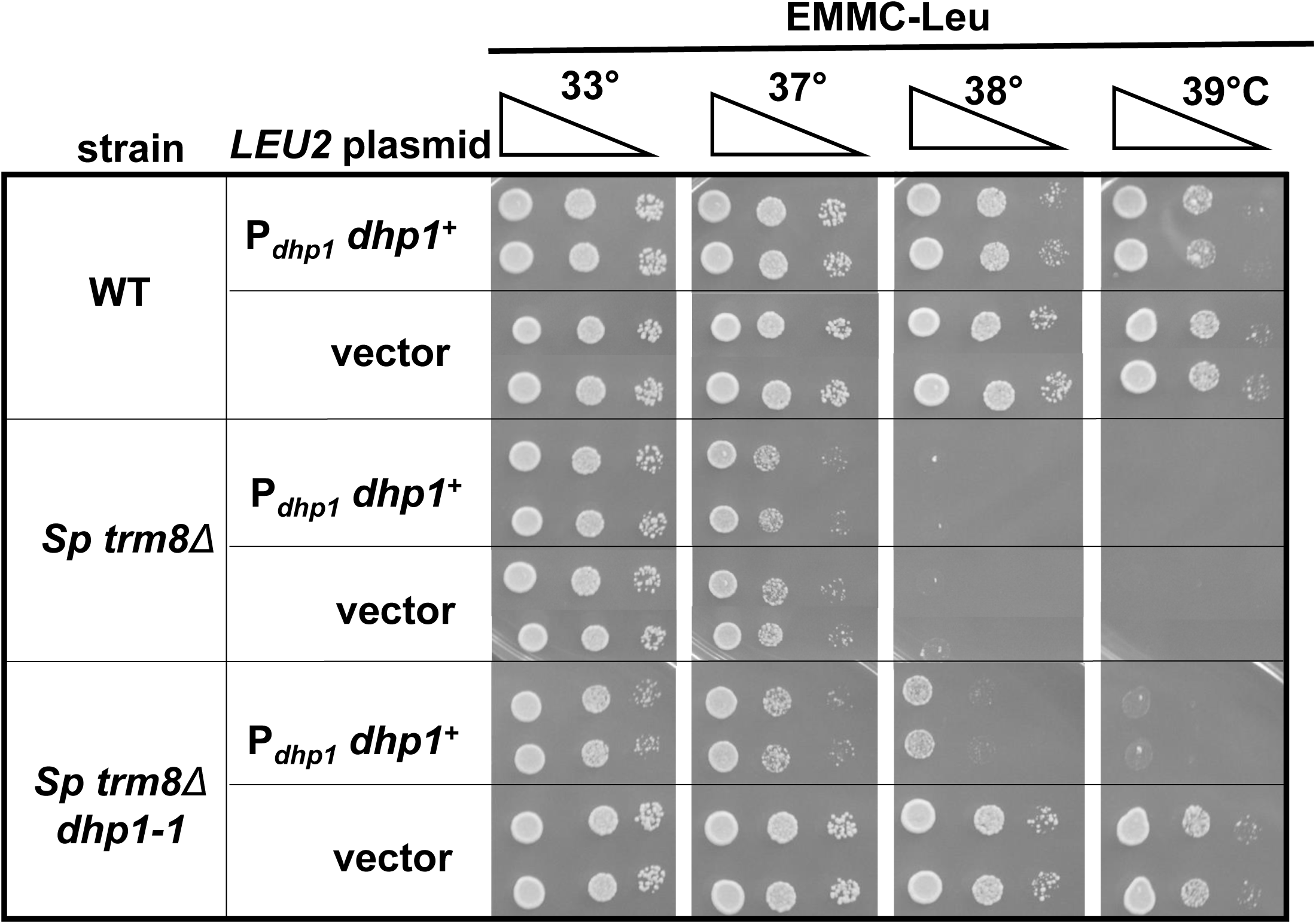
Expression of P*_dhp1_ dhp1^+^* restored temperature sensitive growth in the *S. pombe trm8 dhp1-1* mutant. WT, *trm8Δ*, and *trm8Δ dhp1-1* cells expressing P*_dhp1_ dhp1^+^* or a vector were grown overnight in EMMC-Leu media at 30°C, and analyzed for growth.

**Fig. S10.**
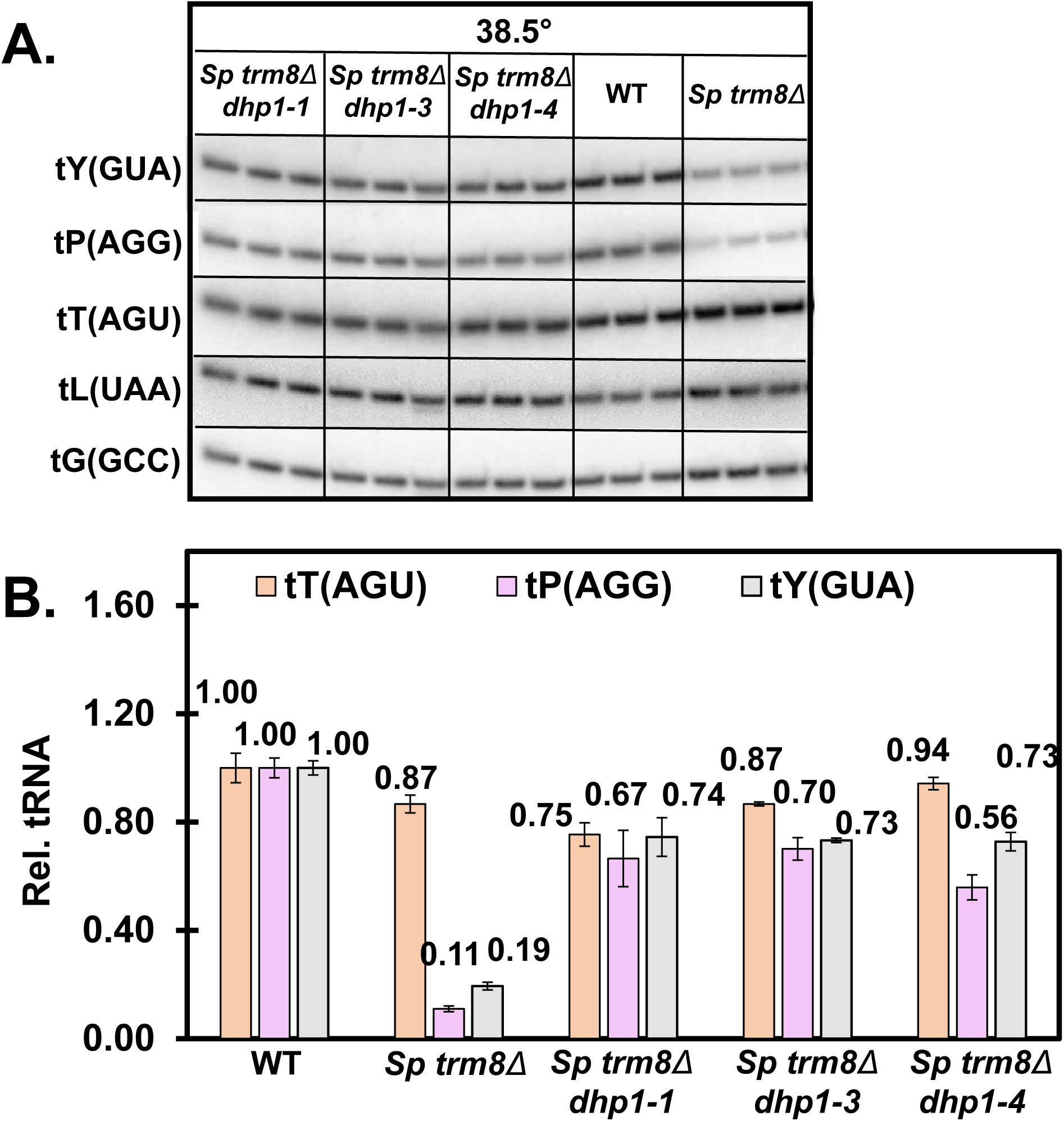
A. *S. pombe trm8Δ dhp1-3* and *trm8Δ dhp1-4* mutants also restored tY(GUA) and tP(AGG) tRNA levels at 38.5 °C. Strains were grown in YES media at 30°C and shifted to 38.5°C for 8 hours, and RNA was isolated and analyzed by Northern blotting as in Fig. 2A. **B. Quantification of tRNA levels in different *S. pombe trm8Δ dhp1* mutants.** tRNA levels were quantified as in Fig. 2B. tT(AGU), brown; tP(AGG), purple; tY(GUA), gray.

**Fig. S11.**
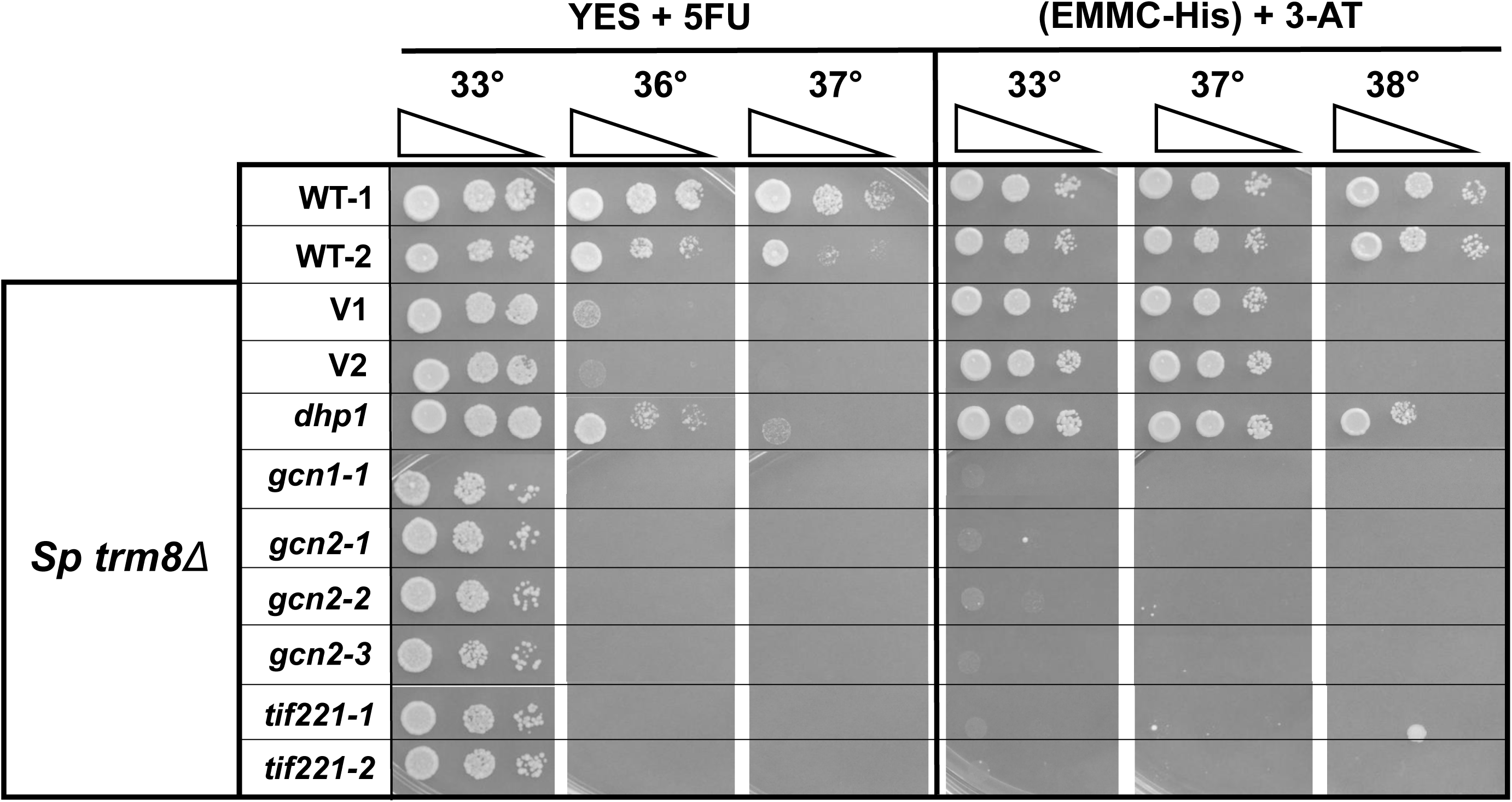
Mutations in the GAAC pathway did not rescue the 5-FU sensitivity of *S. pombe trm8Δ* mutants, and conferred enhanced 3-AT sensitivity. Strains grown for Fig. 4A were analyzed for growth on plates containing YES media + 5-FU (30 µg/ml) and EMMC-His media + 10 mM 3-AT.

**Fig. S12.**
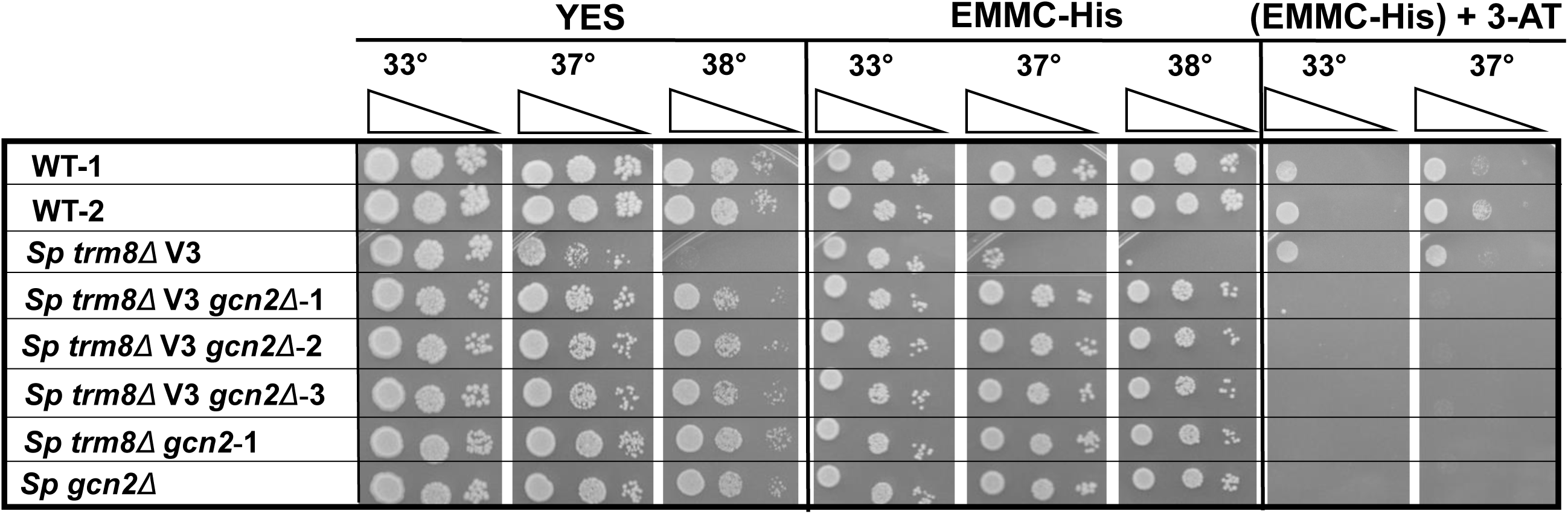
Reconstructed *S. pombe trm8Δ gcn2Δ* mutants had the same growth properties as the original *trm8Δ gcn2-1* strain. Strains were analyzed for growth on YES media, EMMC-His or EMMC-His media containing 10 mM 3-AT, as described in Fig. 1B. Reconstructed *S. pombe trm8Δ* mutant was labeled as V3.

**Fig. S13.**
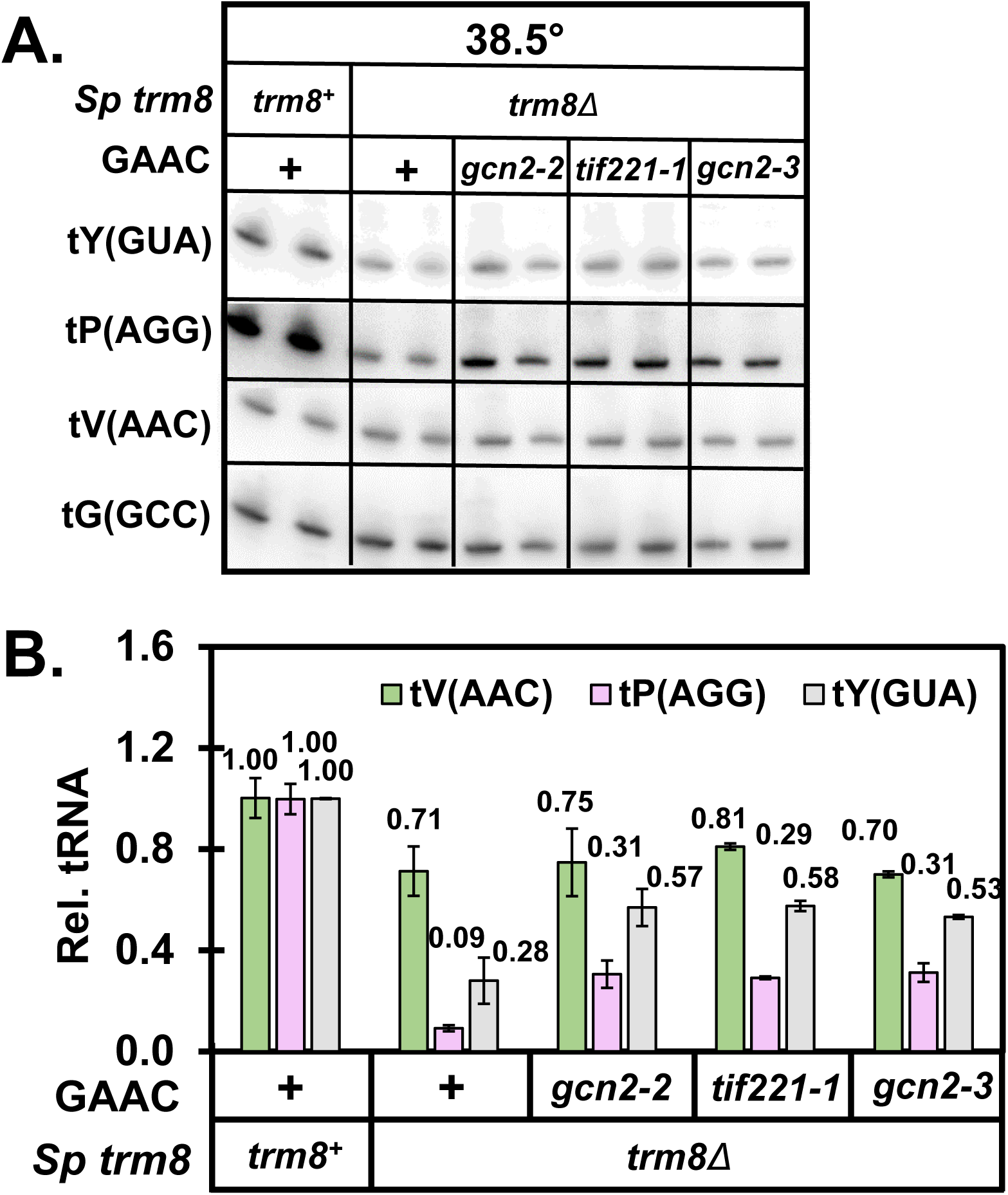
A. Northern analysis of WT, *Sp trm8Δ*, *Sp trm8Δ gcn2-2, trm8Δ tif221-1,* and *trm8Δ gcn2-3* cells. Strains were grown in YES media at 30°C and shifted to 38.5°C for 8 hours, and RNA was isolated and analyzed by Northern blotting as in Fig. 2A. **B. Quantification of tRNA levels.** tRNA levels were quantified as in Fig. 2B. n=2 for all strains.

**Fig. S14.**
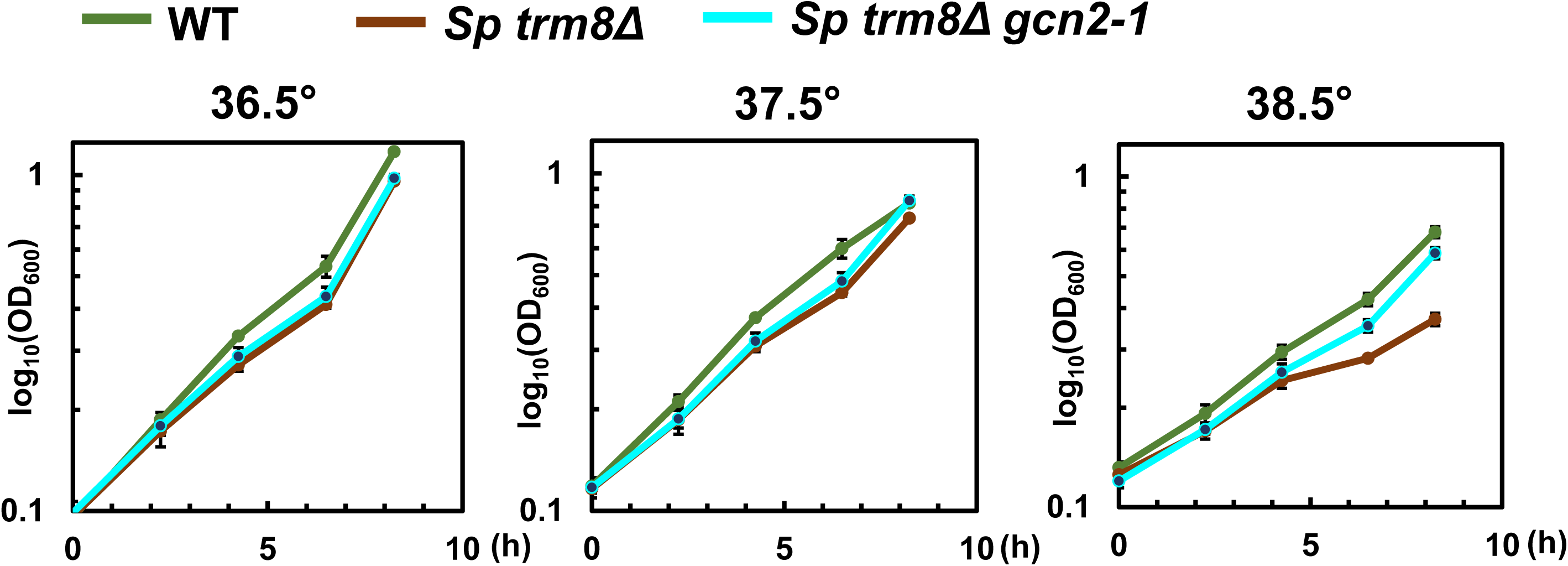
The temperature sensitivity of *S. pombe trm8Δ* mutants observed at 38.5°C in YES liquid media was suppressed in a *trm8Δ gcn2-1* mutant. Strains were grown in YES media at 30°C, shifted to the indicated temperatures, and then growth was monitored for 8 hours before harvest as described in Materials and Methods, and analysis of mRNAs and tRNAs in Fig. 5A, 5B, 5C, and S15.

**Fig. S15.**
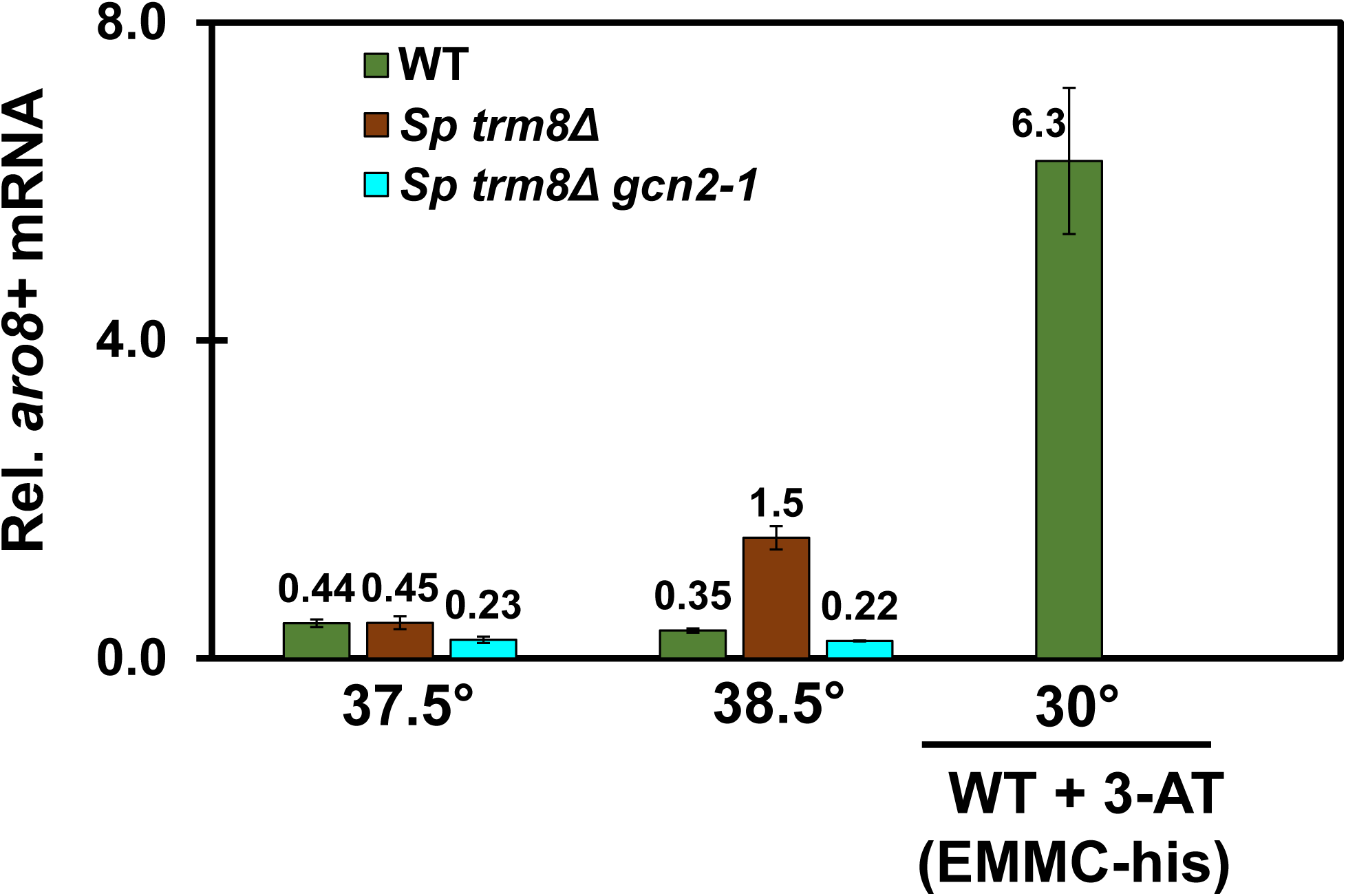
*S. pombe trm8Δ* mutants induced *aro8^+^* mRNA expression at 38.5°C but not at 37.5°C. Bulk RNA from the growth in Fig. S14 was used for the RT-qPCR analysis of *aro8^+^(SPAC56E4.03)* mRNA levels, as in Fig. 5A.

**Fig. S16.**
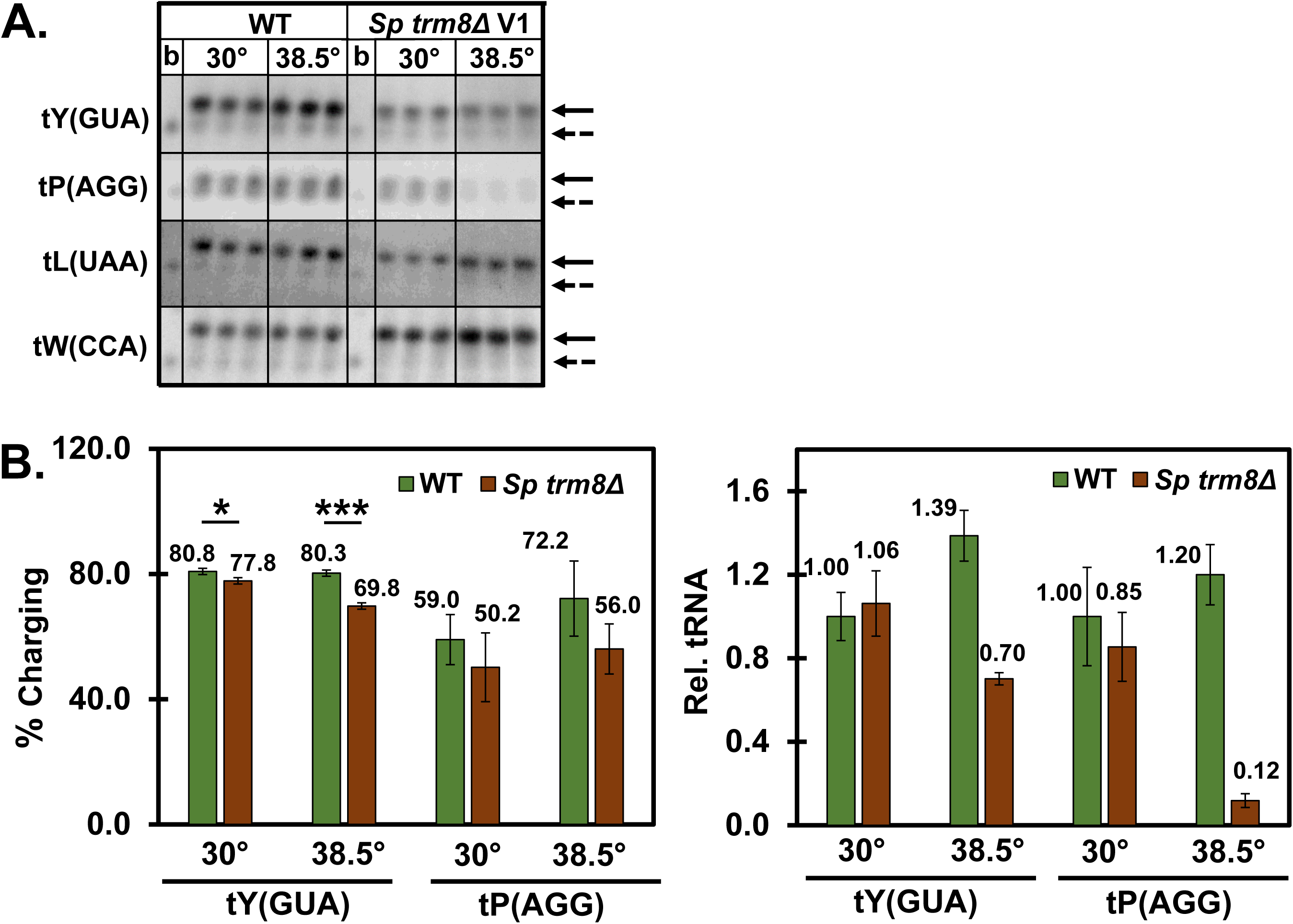
A. Analysis of charging levels of tY(GUA) or tP(AGG) in *S. pombe trm8Δ* mutants at 38.5°C. Strains were grown in YES media at 30°C and shifted to 38.5°C, and samples were harvested after 8 hours. Then bulk RNA was isolated and resolved by denaturing PAGE under acidic conditions (to preserve tRNA charging), transferred, and then analyzed by hybridization as described in Materials and Methods. Control samples (WT and *S. pombe trm8Δ* mutants) were treated with 1 mM EDTA and 0.1 M Tris-HCl (pH 9.0) for 30 min at 37°C to de-acylate the tRNA. b, base treated bulk RNA; Upper arrows, charged tRNA species; lower arrows with dashed lines, uncharged tRNA species**. B. Quantification of tY(GUA) or tP(AGG) charging and tRNA levels.** The percent charging was calculated as the ratio of aminoacylated species to the total for each tRNA. Relative levels of tP(AGG) and tY(GUA) were quantified as in Fig. 2B, relative to the non-Trm8 substrate tL(UAA).

**Fig. S17.**
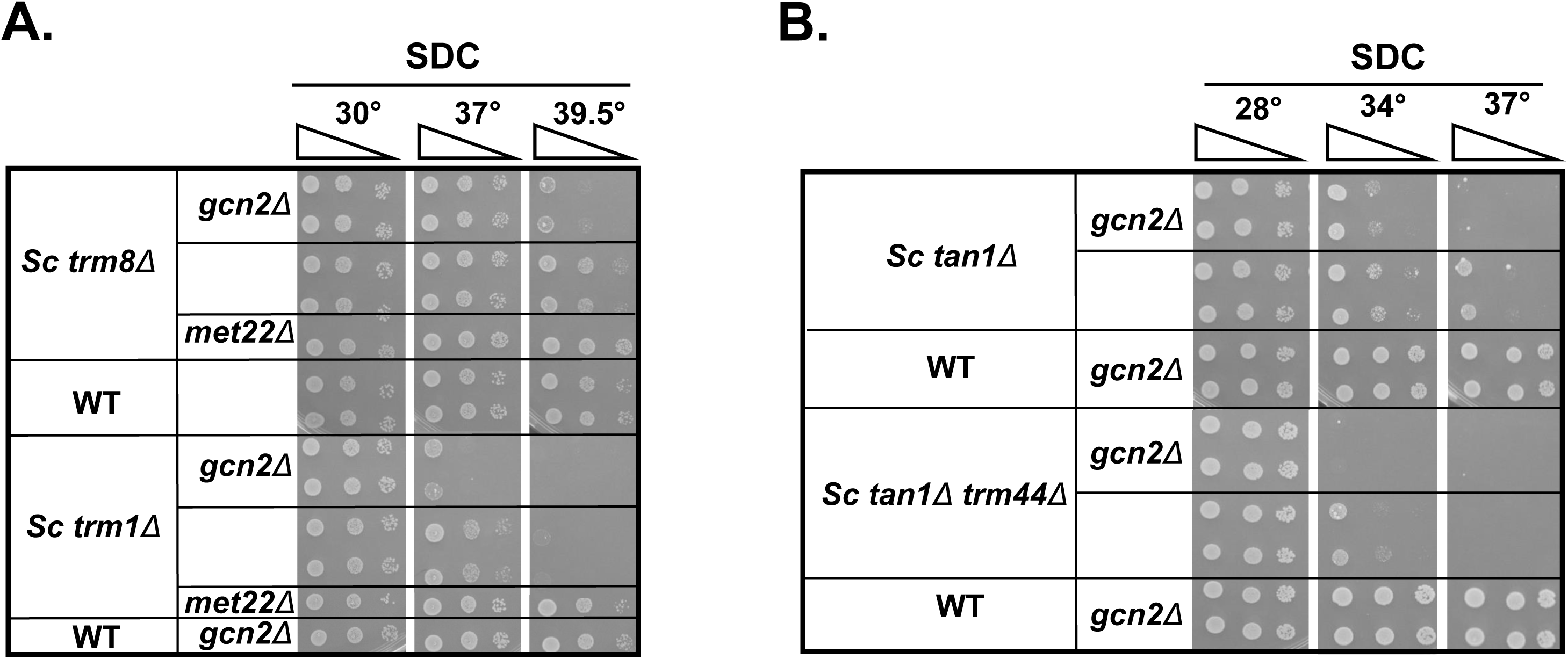
A. Deletion of *GCN2* exacerbated the temperature sensitivity of *S. cerevisiae trm8Δ* and *trm1Δ* mutants in SDC media. Strains were grown overnight in YPD media at 28°C and analyzed for growth on SDC plates. B. Deletion of *GCN2* exacerbated the temperature sensitivity of *S. cerevisiae tan1Δ* and *tan1Δ trm44Δ* mutants in SDC media. Strains were grown overnight in YPD media 28°C and analyzed for growth on SDC plates.

**Fig. S18.**
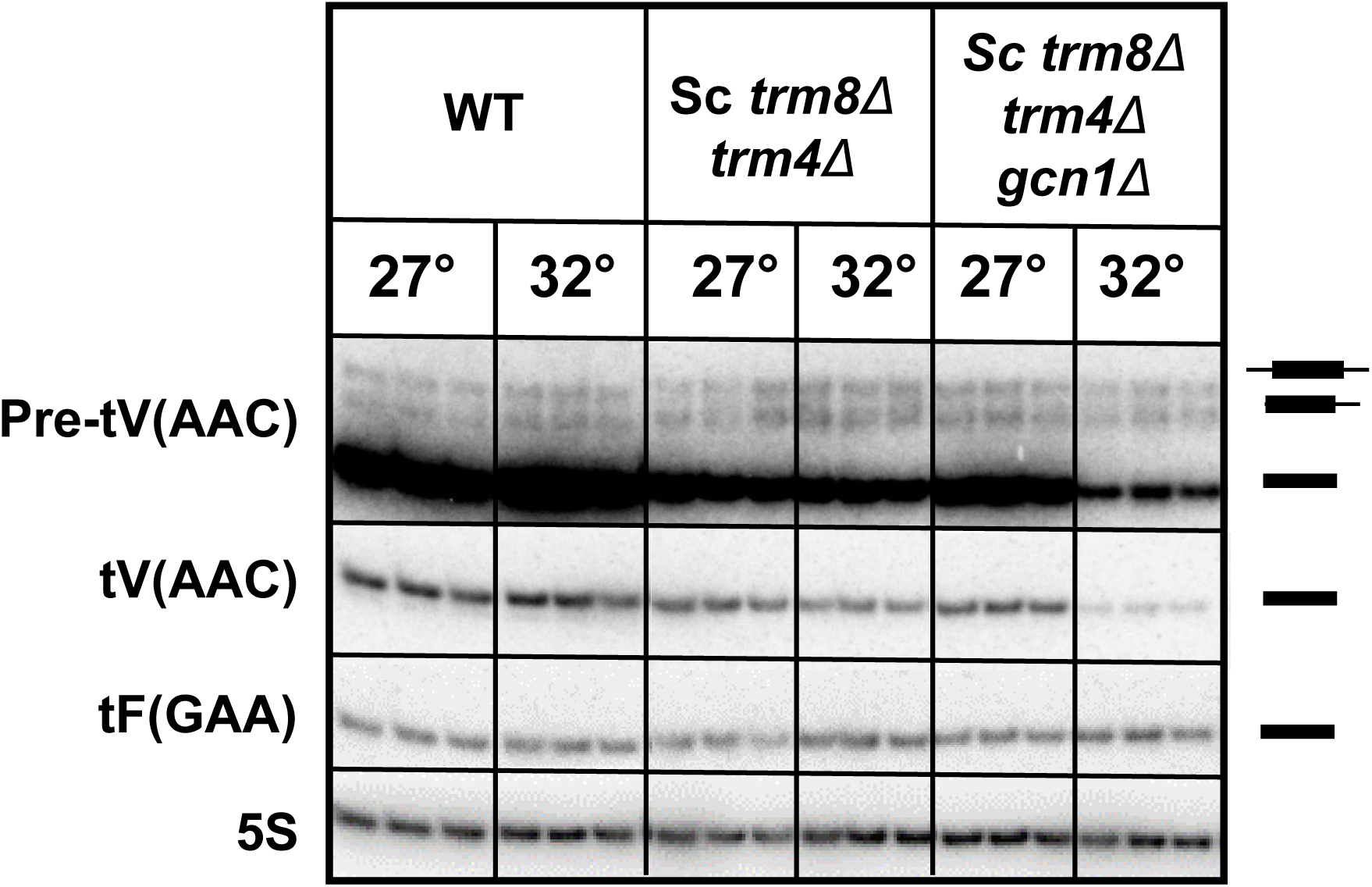
Analysis of pre-tV(AAC) levels in WT, *S. cerevisiae trm8Δ trm4Δ*, and *trm8Δ trm4Δ gcn1Δ* cells after shift from 28°C to 32°C. Bulk RNA from the growth done for Fig. 6B, 6C was used for the northern analysis.

**Fig. S19.**
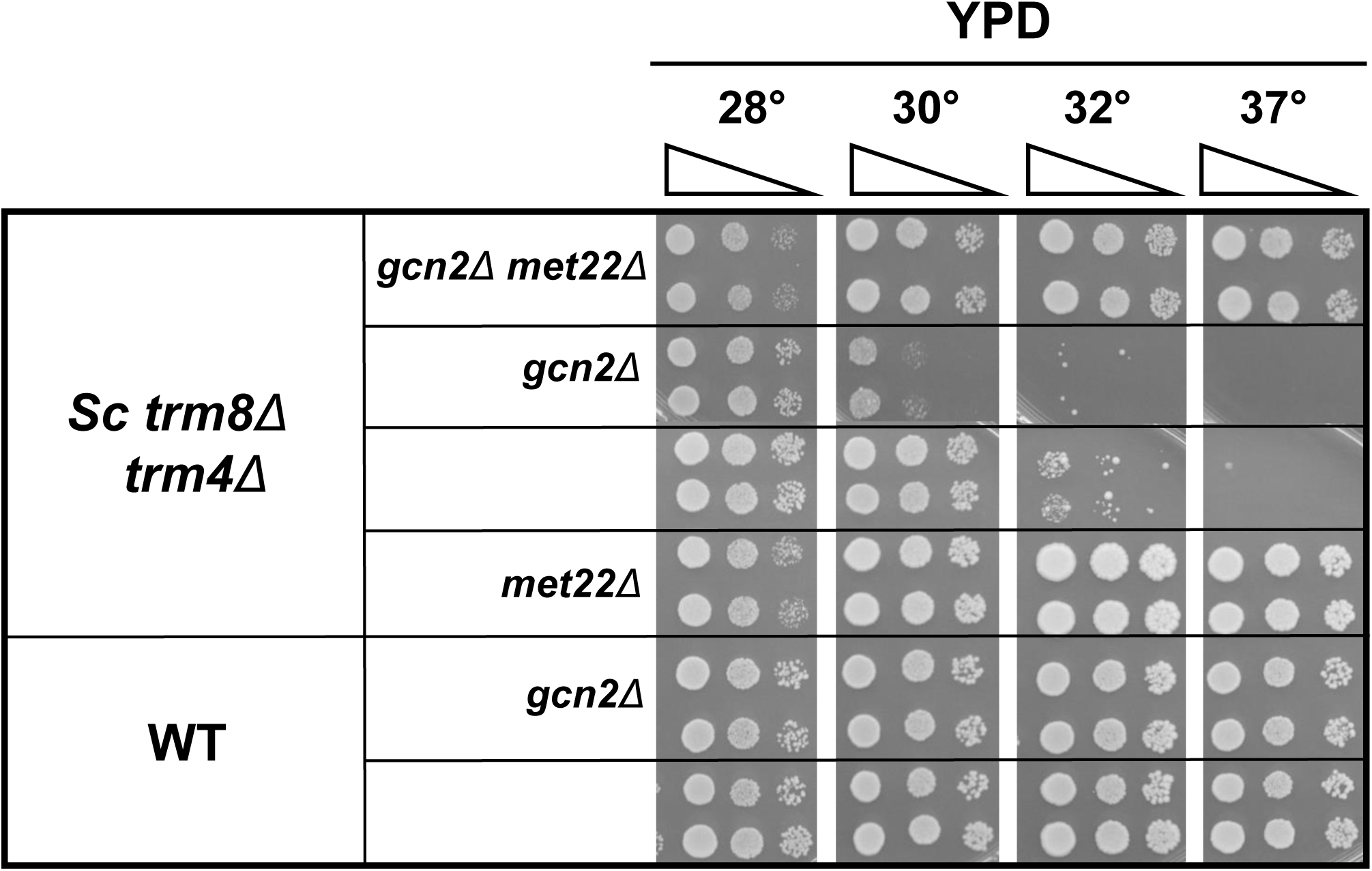
Deletion of *MET22* restored growth of *S. cerevisiae trm8Δ trm4Δ gcn2Δ* mutants at elevated temperatures. Strains were grown overnight in YPD media 28°C and analyzed for growth on YPD plates.

**Figure S20.**
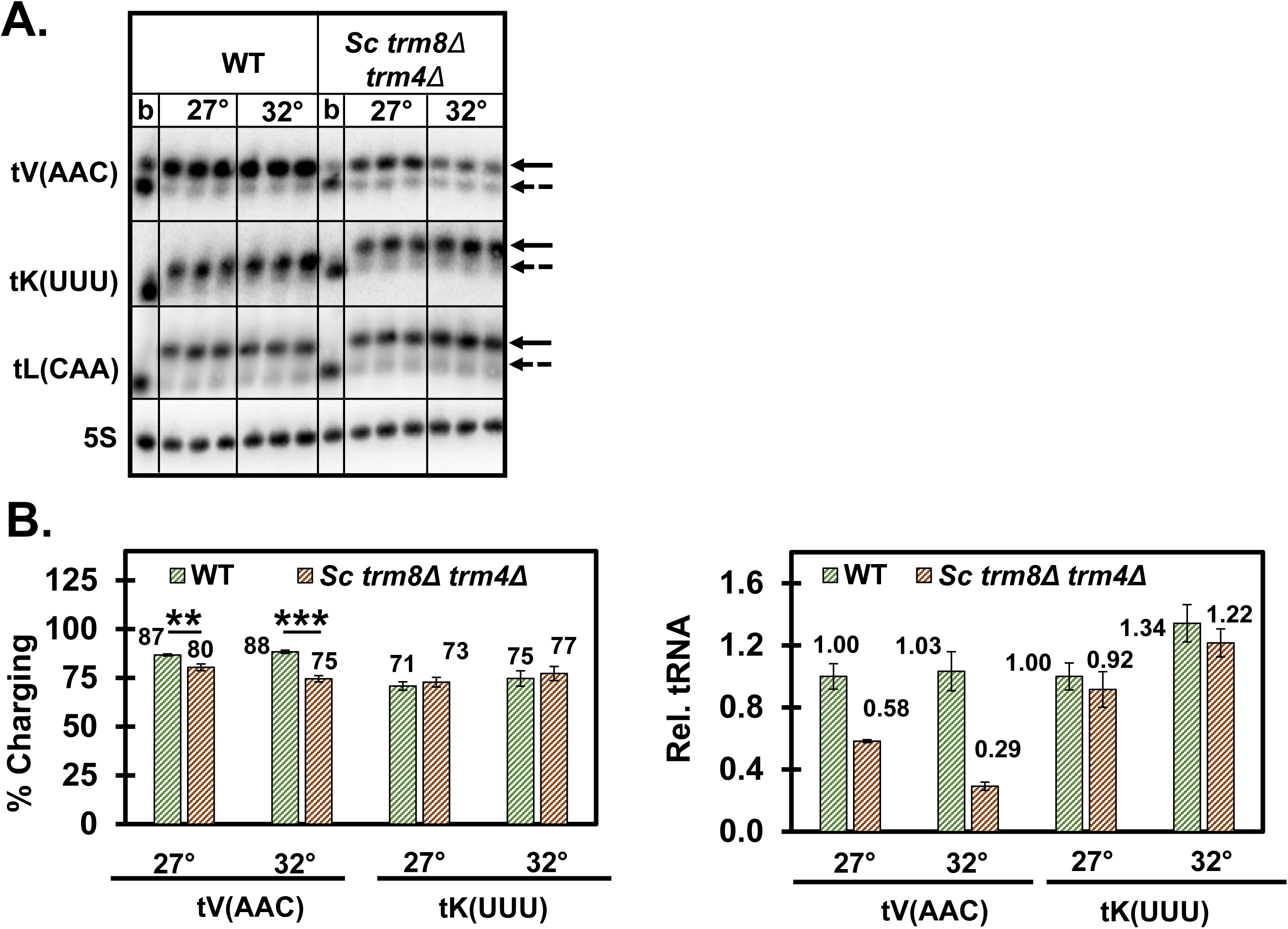
A. Analysis of tRNA charging levels in *S. cerevisiae trm8Δ trm4Δ* mutants after shift to 32°C. Cell pellets from the growth for Fig. 6B were used to isolate acidic RNA and analyzed by acidic northern as described in Fig. S16. **B. Quantification of tRNA charging and tRNA levels.** The percent aminoacylation of tV(AAC) and tK(UUU) was calculated as described in Fig. S16, and relative tRNA levels were quantified as in Fig. 6C.

## Supplementary Tables

**Table S1.**
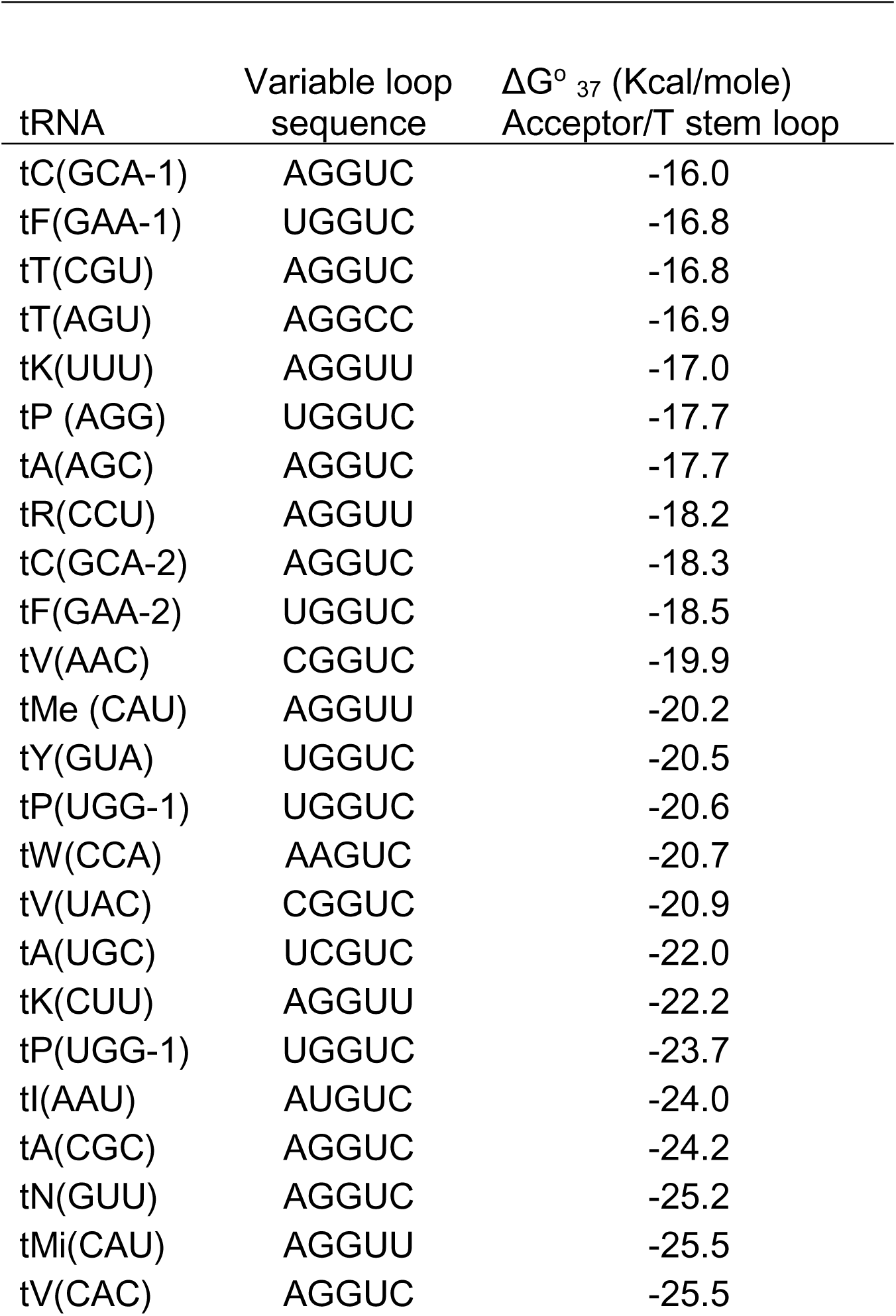
Variable loop sequences and the folding free energies of the acceptor stem/Tstem loop of predicted Trm8 substrate tRNAs of *S. pombe trm8Δ* mutants

**Table S2.** GAAC mutations identified in *S. pombe trm8Δ* suppressors

| Strain | Mutation |
| --- | --- |
| *Sp trm8 gcn1-1* | inframe deletion in V2193-A2195 |
| *Sp trm8 gcn2-1* | start lost, G3A |
| *Sp trm8 gcn2-2* | Frameshift at R71 |
| *Sp trm8 gcn2-3* | Stop at W982 |
| *Sp trm8 tif221-1* | Stop at Q257 |
| *Sp trm8 tif221-2* | Frameshift at L43 |

**Table S3.**
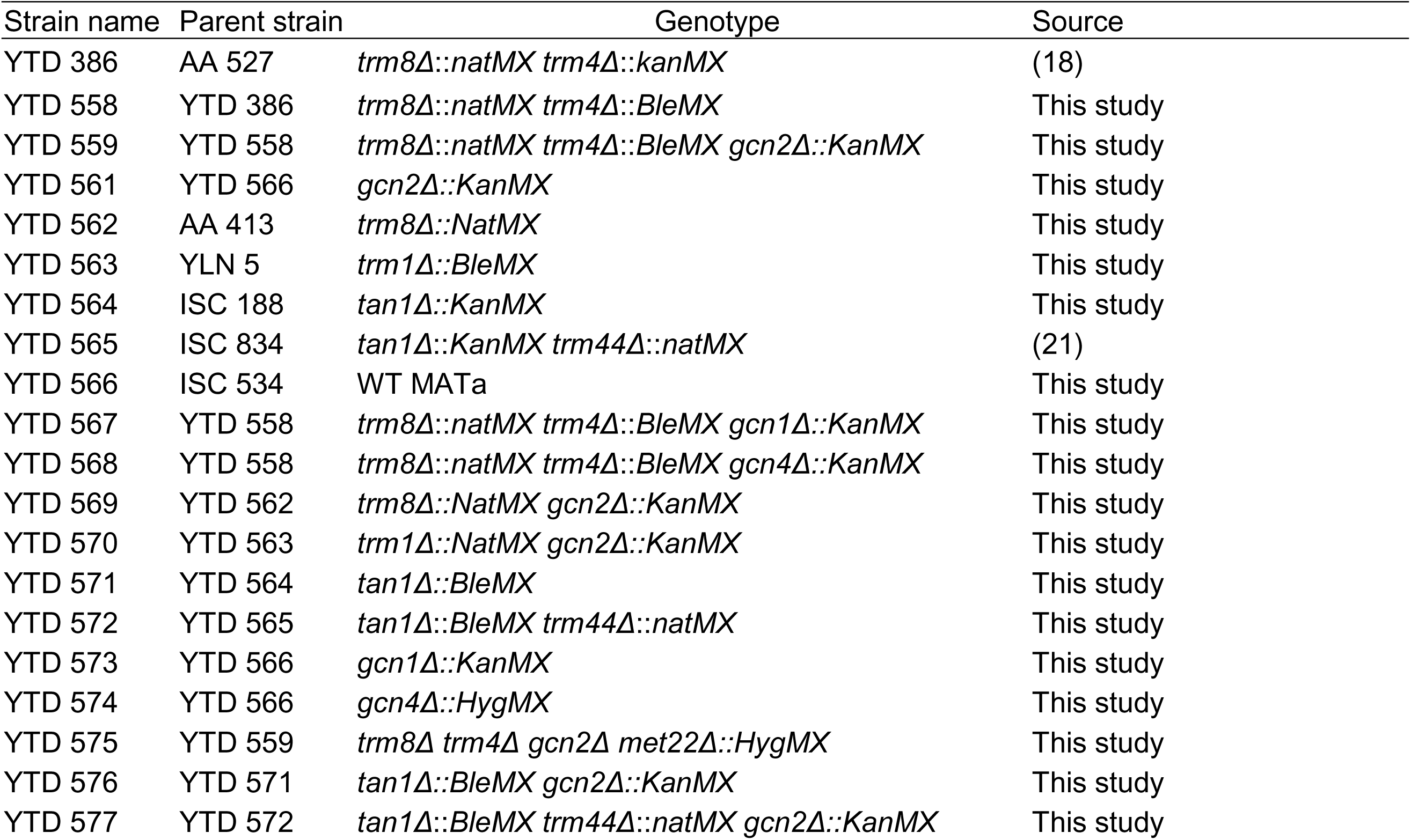
*S. cerevisiae* Strains used in this study

| Strain name | Parent strain | Genotype | Source |
| --- | --- | --- | --- |
| YTD 386 | AA 527 | *trm8Δ::natMX trm4Δ::kanMX* | (18) |
| YTD 558 | YTD 386 | *trm8Δ::natMX trm4Δ::BleMX* | This study |
| YTD 559 | YTD 558 | *trm8Δ::natMX trm4Δ::BleMX gcn2Δ::KanMX* | This study |
| YTD 561 | YTD 566 | *gcn2Δ::KanMX* | This study |
| YTD 562 | AA 413 | *trm8Δ::NatMX* | This study |
| YTD 563 | YLN 5 | *trm1Δ::BleMX* | This study |
| YTD 564 | ISC 188 | *tan1Δ::KanMX* | This study |
| YTD 565 | ISC 834 | *tan1Δ::KanMX trm44Δ::natMX* | (21) |
| YTD 566 | ISC 534 | WT MATa | This study |
| YTD 567 | YTD 558 | *trm8Δ::natMX trm4Δ::BleMX gcn1Δ::KanMX* | This study |
| YTD 568 | YTD 558 | *trm8Δ::natMX trm4Δ::BleMX gcn4Δ::KanMX* | This study |
| YTD 569 | YTD 562 | *trm8Δ::NatMX gcn2Δ::KanMX* | This study |
| YTD 570 | YTD 563 | *trm1Δ::NatMX gcn2Δ::KanMX* | This study |
| YTD 571 | YTD 564 | *tan1Δ::BleMX* | This study |
| YTD 572 | YTD 565 | *tan1Δ::BleMX trm44Δ::natMX* | This study |
| YTD 573 | YTD 566 | *gcn1Δ::KanMX* | This study |
| YTD 574 | YTD 566 | *gcn4Δ::HygMX* | This study |
| YTD 575 | YTD 559 | *trm8Δ trm4Δ gcn2Δ met22Δ::HygMX* | This study |
| YTD 576 | YTD 571 | *tan1Δ::BleMX gcn2Δ::KanMX* | This study |
| YTD 577 | YTD 572 | *tan1Δ::BleMX trm44Δ::natMX gcn2Δ::KanMX* | This study |

**Table S4.** Plasmids used in this study

| Plasmid | Parent | Description | Source |
| --- | --- | --- | --- |
| pREP3X |  | *LEU2 P<sub>nmt1</sub>* | (134) |
| pREP41X |  | *LEU2 P<sub>nmt1\*</sub>* | (134) |
| pREP81X |  | *LEU2 P<sub>nmt1\*\*</sub>* | (134) |
| AB 553-1 | pREP3X | *LEU2* with a NotI between PstI and P<sub>nmt1</sub> | This study |
| ETD 67-1 | AB 553-1 | *LEU2 P<sub>trm8</sub> trm8<sup>+</sup>* | This study |
| ETD 70-1 | AB 553-1 | *LEU2 SPATR<sub>NAPRO.05</sub> SPBTR<sub>NAPRO.05</sub>* | This study |
| ETD 113-1 | AB 553-1 | *LEU2 SPATR<sub>NATYR.01</sub>* | This study |
| ETD 115-1 | AB 553-1 | *LEU2 SPATR<sub>NATYR.01</sub> SPBTR<sub>NAPRO.05</sub>* | This study |
| ETD 104-1 | AB 553-1 | *LEU2 P<sub>dhp1</sub> dhp1<sup>+</sup>* | This study |
| ETD 109-1 | pREP81X | *LEU2 P<sub>nmt1\*\*\*</sub> gcn2<sup>+</sup>* | This study |
| ETD 119-1 | pREP41X | *LEU2 P<sub>nmt1\*\*</sub> tif221<sup>+</sup>* | This study |
| ETD 126 | EIC-17 | tV(AAC)E2 URA3 2u | (39) |

**Table S5.**
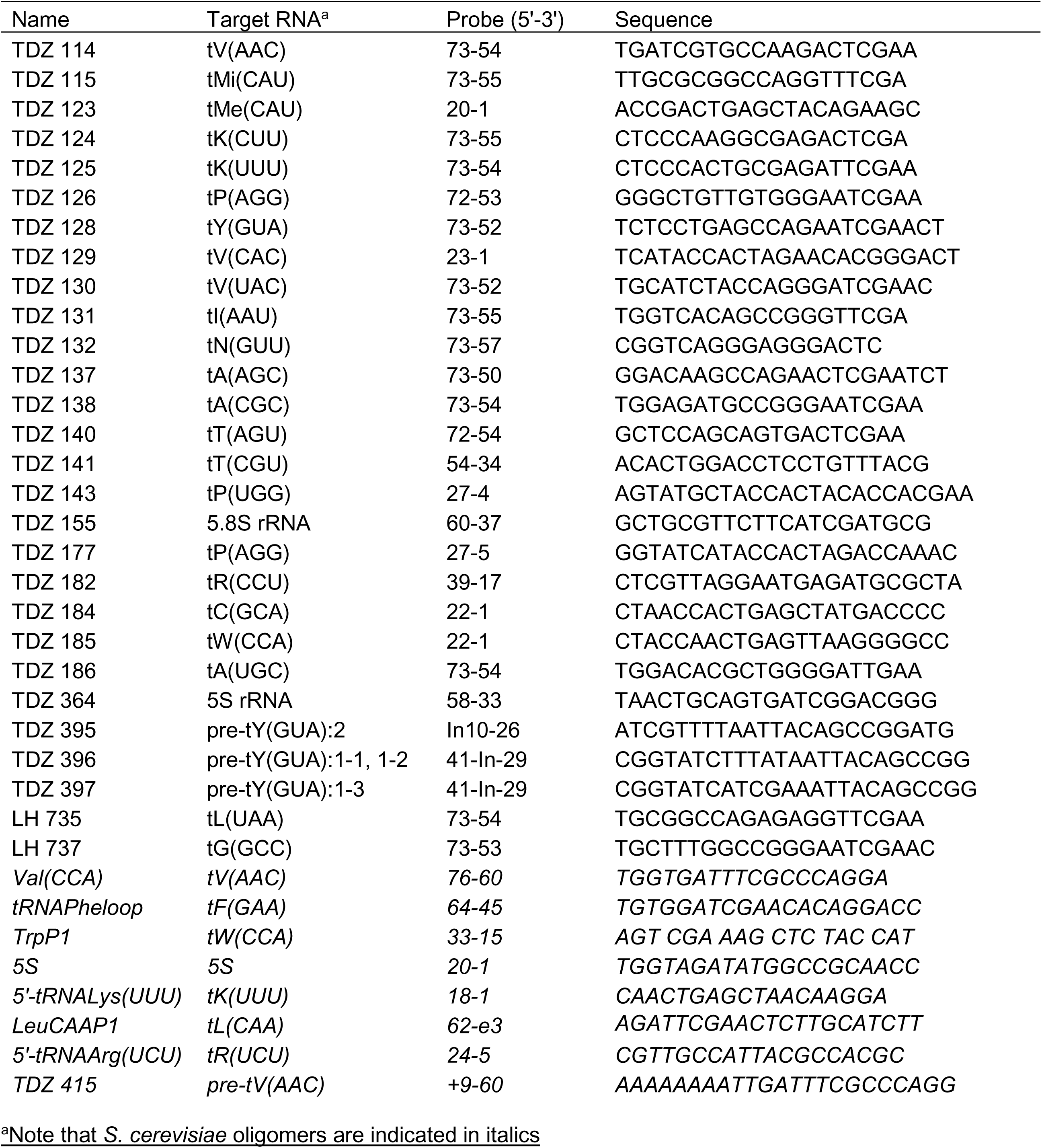
Oligomers used for northern analysis

**Table S6.**
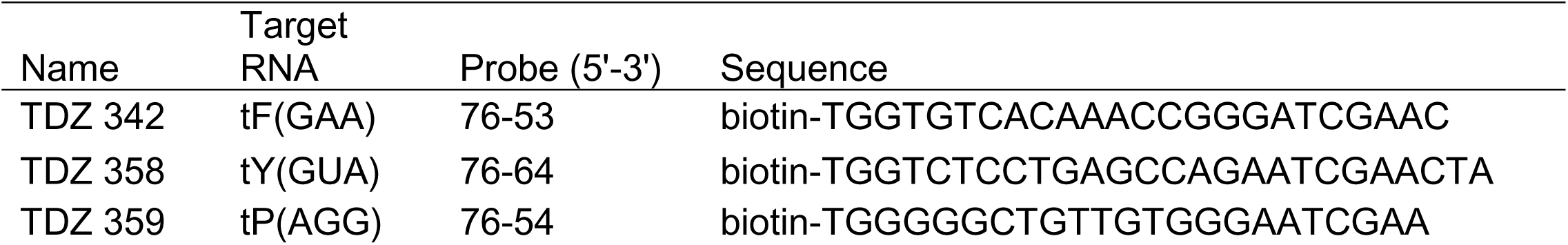
Oligomers used for tRNA purifications

